# Stable and robust Xi and Y transcriptomes drive cell-type-specific autosomal and Xa responses *in vivo* and *in vitro* in four human cell types

**DOI:** 10.1101/2024.03.18.585578

**Authors:** Laura V. Blanton, Adrianna K. San Roman, Geryl Wood, Ashley Buscetta, Nicole Banks, Helen Skaletsky, Alexander K. Godfrey, Thao T. Pham, Jennifer F. Hughes, Laura G. Brown, Paul Kruszka, Angela E. Lin, Daniel L. Kastner, Maximilian Muenke, David C. Page

## Abstract

Recent *in vitro* studies of human sex chromosome aneuploidy showed that the Xi (“inactive” X) and Y chromosomes broadly modulate autosomal and Xa (“active” X) gene expression in two cell types. We tested these findings *in vivo* in two additional cell types. Using linear modeling in CD4+ T cells and monocytes from individuals with one to three X chromosomes and zero to two Y chromosomes, we identified 82 sex-chromosomal and 344 autosomal genes whose expression changed significantly with Xi and/or Y dosage *in vivo*. Changes in sex-chromosomal expression were remarkably constant *in vivo* and *in vitro* across all four cell types examined. In contrast, autosomal responses to Xi and/or Y dosage were largely cell-type-specific, with up to 2.6-fold more variation than sex-chromosomal responses. Targets of the X- and Y-encoded transcription factors ZFX and ZFY accounted for a significant fraction of these autosomal responses both *in vivo* and *in vitro*. We conclude that the human Xi and Y transcriptomes are surprisingly robust and stable across the four cell types examined, yet they modulate autosomal and Xa genes – and cell function – in a cell-type-specific fashion. These emerging principles offer a foundation for exploring the wide-ranging regulatory roles of the sex chromosomes across the human body.

## Introduction

The somatic cells of human females and males share forty-five chromosomes: 22 pairs of autosomes and one X chromosome in the epigenetically ‘active’ state (Xa). The sexes are distinguished by the forty-sixth chromosome, which is either an ‘inactive’ X chromosome (Xi) in somatic cells of most females or a Y chromosome (Chr Y) in males.^1,2^ Despite the ‘inactive’ label, Xi is transcriptionally active, and contributes substantially to female fetal viability.^3–5^ In the reproductive tract, the male-differentiating roles of Chr Y are well documented.^6–10^ Elsewhere in the human body, we know little about Xi and Y’s roles in gene expression and its regulation genome-wide.

We recently assessed how Xi and Y chromosomes impact gene expression across the genome by RNA-sequencing of lymphoblastoid cell lines (LCLs) and fibroblasts cultured from individuals with sex chromosome aneuploidy (one to four X chromosomes and zero to four Y chromosomes).^11,12^ Through linear modeling, we quantified the effects of Xi- and Y-chromosome dosage on sex-chromosomal^11^ and autosomal^12^ gene expression. These studies also identified zinc finger transcription factors encoded by the homologous X-Y gene pair *ZFX* and *ZFY* as major drivers of the autosomal gene response.^12^ In these studies, the highly controlled setting of *in vitro* culture was both a strength and a limitation: a strength in that the effects of circulating hormones and environmental exposures were minimized, and a limitation in that the pertinence of the findings to human cells *in vivo* remained unknown.

To address this limitation of our previous studies, we assembled parallel collections of primary CD4+ T cells and monocytes, representing the lymphoid and myeloid arms of the immune system, isolated from sex chromosome euploid and aneuploid individuals. We prioritized study of these peripheral immune cells for two reasons. First, they are among the most accessible primary cells in humans. Second, sex differences within the immune system are pervasive in health and disease. Across hematopoietic lineages, nearly every major immune cell type – from B and T cells in the adaptive immune arm to monocytes and neutrophils in the innate immune arm – displays sex-biased phenotypes.^13–17^ Immunological sex differences profoundly impact human health, with disparities in the prevalence and presentation of autoimmunity and distinct responses to infections and vaccines.^18–20^ Even in healthy basal conditions, XX and XY individuals show different proportions of circulating immune cells and different immune cell gene expression profiles across the genome.^21–24^

To quantitatively assess Xi and Y dosage effects *in vivo*, we RNA-sequenced CD4+ T cells and monocytes from individuals with one, two, or three X chromosomes and zero, one, or two Y chromosomes, constructing linear models of both sex-chromosomal and autosomal gene expression responses to increasing Xi and Y chromosome dosage. As reported here, we found the Xi and Y transcriptomes to be remarkably modular, stable, and robust – not only between the two *in vivo* cell types but also when compared with the previously studied *in vitro* LCLs and fibroblasts. In contrast, the effects of Xi and Y dosage on both autosomal and Xa expression were significantly more variable across cell types. These findings indicate that a consistent set of genes expressed on Xi and Y chromosomes drive cell-type-dependent autosomal and Xa responses, with important implications for understanding the molecular mechanisms that underpin sex-biased phenotypes.

## Results

### Responses of sex-chromosomal genes to sex chromosome dosage in CD4+ T cells and monocytes

To investigate the contribution of sex chromosome constitution to immune cell gene expression, we isolated and performed RNA-Seq on CD4+ T cells and monocytes from, respectively, 76 and 72 adults, with one of six different sex chromosome constitutions, comprising one, two, or three X chromosomes together with zero, one, or two Y chromosomes (**Figure 1A, B; Table S1**). The individuals ranged in age from 17 to 70 years, with a median age of 34.5 years. Computational deconvolution of the bulk RNA-Seq data confirmed successful enrichment of either CD4+ T cells (estimated purity 80.9 +/− 6.7%) or monocytes (81.5 +/− 5.8%; **Figure S1**).^25^

**Figure 1.**
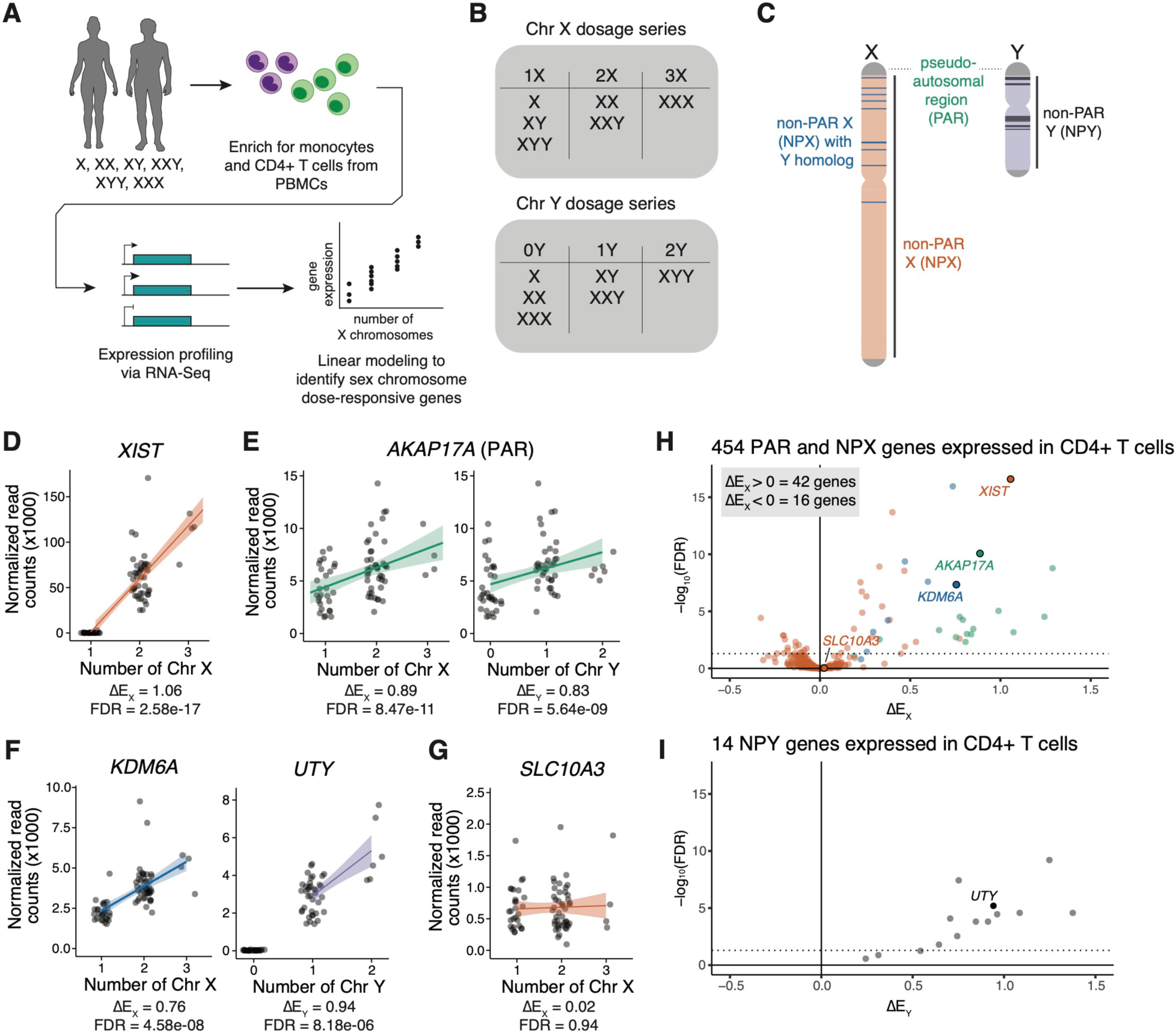
Changes in Xi and Chr Y dosage alter expression of X- and Y-chromosomal genes. **(A)** Experimental design for analysis of primary CD4+ T cells and monocytes from individuals with varying numbers of sex chromosomes. **(B)** Chr X and Chr Y dosage series in CD4+ T cells and monocytes. **(C)** Schematic of Chr X and Chr Y, indicating locations of pseudoautosomal regions (PAR) in green, non-PAR X (NPX) in orange, NPX genes with NPY homologs in blue, and NPY genes in black. **(D - G)** Normalized read counts (x1000) by sex chromosome dosage for *XIST* **(D)**, the PAR gene *AKAP17A* **(E)**, the NPX-NPY homologous pair *KDM6A* and *UTY* **(F)**, or the non-Xi- or Chr Y-responsive *SLC10A3* **(G)** in CD4+ T cells. Regression lines with confidence intervals are shown. **(H, I)** Volcano plot of ΔE_X_ values for all expressed NPX and PAR genes **(H)** or ΔE_Y_ values of all expressed NPY genes in CD4+ T cells. Dotted horizontal lines indicate adjusted *p* < 0.05. In **(H)**, genes are annotated by class: PAR genes in green, NPX genes with NPY homologs in blue, and NPX genes with no NPY homolog in orange.

We first examined gene expression from Chr X and Chr Y, including genes in the pseudoautosomal region (PAR), the region of sequence identity and meiotic crossing-over between Chr X and Chr Y.^26^ After filtering for genes with at least one transcript per million (TPM) in either euploid karyotype, there were 18 PAR genes expressed in CD4+ T cells and 19 in monocytes; 436 non-PAR X-chromosomal (NPX) genes expressed in CD4+ T cells and 390 in monocytes; and 14 non-PAR Y-chromosomal (NPY) genes expressed in CD4+ T cells and 13 in monocytes. As in our prior studies of LCLs and fibroblasts, we used a linear model to assess the modular impact of Chr X or Chr Y dosage on a given gene’s expression, controlling for sequencing batch; donor age had little impact on the model and was excluded from downstream analyses (**Figure S2**). To quantify the relative expression of a given gene in the presence of additional copies of Xi or Chr Y, we employed the metrics ΔE_X_ or ΔE_Y_, respectively, by dividing the slope of regression (β_X_ or β_Y,_ representing the change in expression per copy of Chr X or Y) by the intercept (β_0,_ representing expression in samples with one Chr X or Y). From these ΔE_X_ and ΔE_Y_ values, we identified genes whose expression significantly increased with additional copies of Xi or Chr Y (ΔE_X_ or ΔE_Y_ > 0), significantly decreased (ΔE_X_ or ΔE_Y_ < 0), or remained essentially unchanged (ΔE_X_ or ΔE_Y_ ≅ 0).

The evolution of the human X and Y chromosomes from ordinary autosomes has led to three distinct classes of X-chromosomal genes expressed in immune and other somatic cells. These include 1) PAR genes, 2) NPX genes with diverged homologs on Chr Y, and 3) NPX genes that have lost their homologs on Chr Y (**Figure 1C**).^10,11^ Fifteen of the 18 PAR genes expressed in CD4+ T cells and 17 of the 19 PAR genes expressed in monocytes significantly increased expression with Xi dosage, while 10 PAR genes in CD4+ T cells and 15 in monocytes significantly increased expression with Chr Y dosage. Forty-three NPX and 11 NPY genes responded significantly to, respectively, Xi or Chr Y dosage in CD4+ T cells, and 17 NPX and nine NPY genes responded significantly to Xi or Y dosage in monocytes (**Table S2, Table S3**). Of the 43 NPX genes that responded to Xi dosage in CD4+ T cells, seven possessed an NPY homolog; of the 17 NPX genes that responded to Xi dosage in monocytes, six possessed an NPY homolog. As expected, the *XIST* lincRNA, which mediates X chromosome inactivation, was expressed in individuals with at least two X chromosomes, and its expression increased with additional Xi chromosomes (**Figure 1D; Figure S3A)**. Genes previously shown to be expressed on both Xa and Xi showed significant positive ΔE_X_ values, including PAR genes and NPX genes with NPY homologs (**Figure 1E-F, 1H; Figure S3B, C**).^5,27^ In contrast, most expressed NPX genes lacking NPY homologs had ΔE_X_ values near zero, indicating little or no change in expression with increasing Xi dosage (**Figure 1G; Figure S3D, E, G, H**). These non-responsive NPX genes included *TLR7* and *CD40LG*, which have roles in immune cell signaling and have been proposed as drivers of female-biased immune phenotypes, but did not change significantly with additional X chromosomes (*TLR7* ΔE_X_ = 0.04 ± 0.12 in monocytes; *CD40LG* ΔE_X_ = - 0.003± 0.04 in CD4+ T cells; **Figure S3D**).^28–32^ Most expressed NPY genes had significant positive ΔE_Y_ values of at least 0.5, with essentially no ΔE_Y_ values near zero (**Figure 1I, Figure S3F**). Integrating these results, we find that ∼95% of expressed PAR genes, 80% of expressed NPX genes with an NPY homolog, ∼8.6% of expressed NPX genes with no NPY homolog, and ∼86% of expressed NPY genes demonstrate robust responses to Xi and/or Y dosage in CD4+ T cells and/or monocytes *in vivo* (**Tables S2, S3**).

### Responses of X-chromosomal genes to Xi dosage are highly similar across four cell types

To understand the responses of X-chromosomal genes to Xi dosage across cell types, we first examined ΔE_X_ and ΔE_Y_ values by gene category: PAR, NPX with NPY homologs, and NPX with no NPY homolog. Regardless of cell type, expressed PAR genes displayed strongly positive values for both ΔE_X_ and ΔE_Y_ (across four cell types, mean ΔE_X_ = 0.78 +/− 0.35 and mean ΔE_Y_ = 0.81 +/− 0.47; **Figure 2A, B**). While expressed NPX genes with NPY homologs also displayed significant positive responses to Xi dosage in each cell type, their ΔE_X_ values were lower than those of expressed PAR genes (mean = 0.36 +/− 0.23), and they showed little or no response to Chr Y dosage. ΔE_X_ values for expressed NPX genes with no NPY homologs were largely near zero, regardless of cell type (mean = 0.03 +/− 0.3) (**Figure 2B**). These findings agree with prior expectations: PAR genes are identical on Chrs X and Y, and each additional Xi or Y chromosome adds an increment of PAR gene expression comparable to that of Xa. Expressed NPX genes with NPY homologs are transcribed on both Xa and Xi in somatic cells (contributing to their positive ΔE_X_ values), but their expression on Xi is attenuated compared to that on Xa. These patterns were evident regardless of whether the cells had been isolated directly from blood (the primary immune cells) or cultured *in vitro* (the previously collected LCLs and fibroblasts) (**Figure 2B-C**).^11^

**Figure 2.**
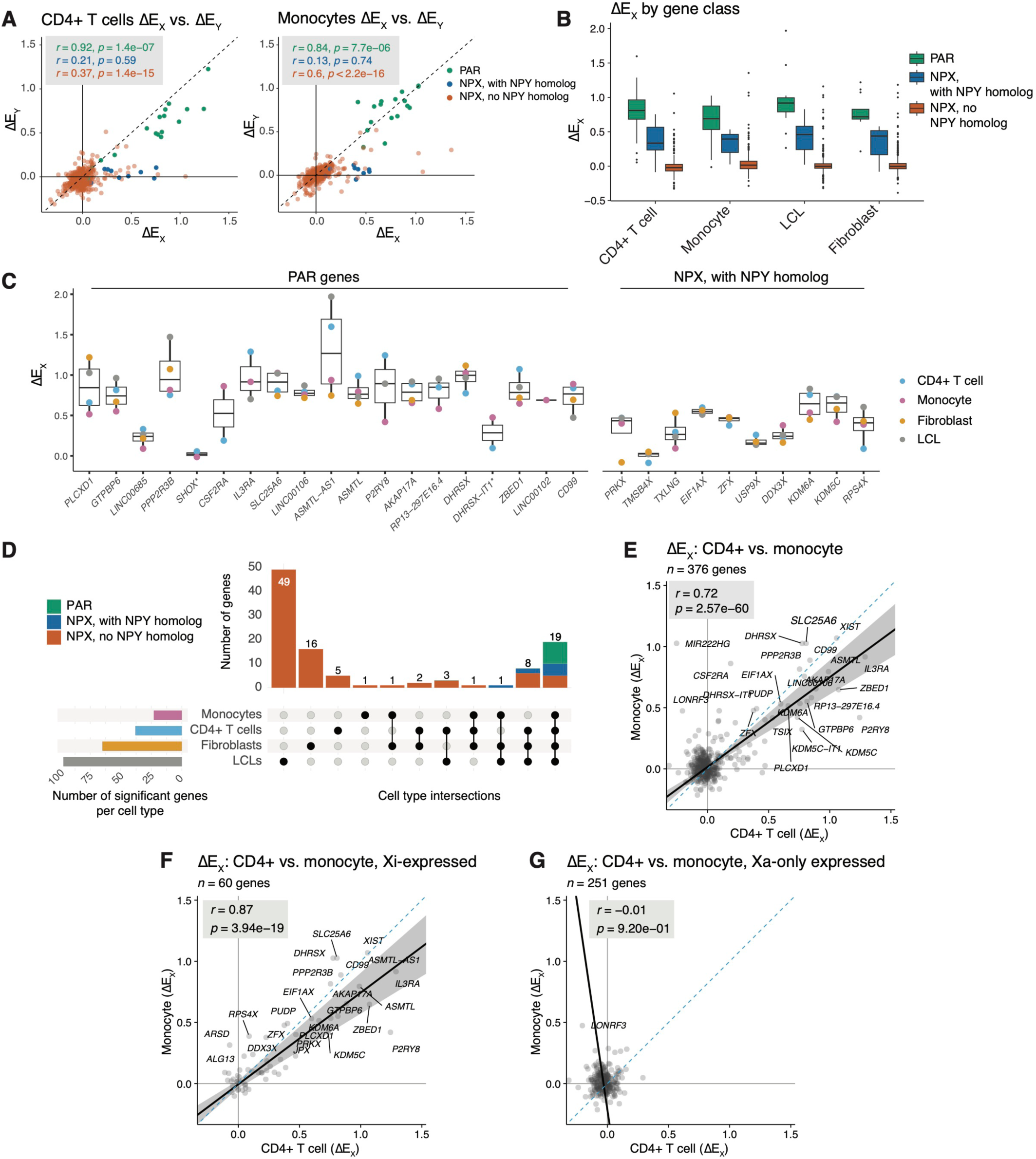
Responses of X-chromosomal genes to Xi and Chr Y dosage across gene classes and cell types. **(A)** Scatter plots of ΔE_X_ versus ΔE_Y_ values of X-chromosomal genes, colored by gene class, in CD4+ T cells and monocytes. Pearson correlation coefficients shown; colored by corresponding gene class. Dashed line indicates X=Y identity line. **(B)** Box plots showing median (line), interquartile range (IQR; top and bottom of box), and 1.5 * IQR (whiskers) of ΔE_X_ values for X-chromosomal genes by cell type and gene class. **(C)** Box plots showing median (line), interquartile range (IQR; top and bottom of box), and 1.5 * IQR (whiskers) of ΔE_X_ values for individual PAR and NPX genes with NPY homologs. Asterisks mark two genes with < 3 TPM median expression in each cell type. **(D)** UpSet plot of genes with statistically significant ΔE_X_ values across CD4+ T cells, monocytes, LCLs, and fibroblasts; stacked bar plots colored by PAR genes in green, NPX genes in orange, NPX genes with NPY homologs in blue. **(E)** Scatter plot of ΔE_X_ values for all expressed X-chromosomal genes in CD4+ T cells and monocytes. Pearson correlation coefficient shown. **(F, G)** Scatter plot of ΔE_X_ values for genes previously annotated as expressed on Xi (“Xi-expressed”) **(F)** or not expressed from Xi (“Xa-only expressed”) **(G)** in CD4+ T cells and monocytes. Pearson correlation coefficients shown.

We next looked across cell types at the genes significantly responsive to Xi dosage. Expressed PAR genes and expressed NPX genes with NPY homologs consistently showed significant responses to Xi dosage across all four cell types. Genes that showed significant responses in only one or two cell types were in all instances NPX genes with no NPY homolog (**Figure 2D**). Fewer X-chromosomal genes reached statistical significance *in vivo* compared to *in vitro*, which is likely driven by multiple factors: the *in vivo* data are subject to uncontrolled environmental variables as the cells are isolated directly from blood, and our *in vivo* dataset has a more limited range of sex chromosome constitutions and fewer total samples. Sub-sampling analysis revealed that our identification of significantly responsive X-chromosomal genes was approaching saturation, with the majority of significantly responsive genes identified but additional genes likely to be identified with additional samples (**Figure S4A, B**). In all, we see that expressed PAR genes and NPX genes with NPY homologs respond more robustly and reliably to Xi dosage than NPX genes lacking an NPY homolog.

Finally, we compared each X-chromosomal gene’s ΔE_X_ values across the four cell types to see how those values were affected by somatic cell differentiation. Although fewer X-chromosomal genes displayed significantly non-zero ΔE_X_ values in the cells collected *in vivo* than in cultured LCLs and fibroblasts, the ΔE_X_ values *in vivo* and *in vitro* were highly correlated (Pearson *r* range was 0.70 – 0.86, all *p* < 2.2e-16, **Figure 2E; Figure S5**). Cognizant that ΔE_X_ reflects both gene expression on Xi and modulation of gene expression on Xa, we divided Chr X genes into those previously shown to be expressed on Xi (and on Xa) or silenced on Xi (expressed only on Xa) (as tallied in Table S4 in San Roman, *et al.*, 2024).^5,11,27,33–36^ This parsing allowed us to see that the consistency of ΔE_X_ values across cell types was a feature of Xi-expressed genes but not of Xi-silenced genes (“Xa-only expressed”) (**Figure 2F, G; Figure S5**).

While nearly all genes on Xi are silenced or show attenuated expression, genes on supernumerary Y chromosomes display no such attenuation of their expression.^11,12^ Examining the impact of additional Y chromosomes *in vivo*, we found that ΔE_Y_ values for NPY genes do not vary as widely as ΔE_X_ values for NPX genes do; rather, median ΔE_Y_ values across the NPY genes within each cell type were largely consistent, ranging from 0.65 to 1.07 among broadly expressed NPY genes (**Figure S6**). These results highlight the distinct effects of adding Xi versus Chr Y to the cell, as supernumerary Y chromosomes show no evidence of dosage compensation through Y inactivation or other mechanisms.

### Autosomal responses to sex chromosome dosage in vivo

Having established the direct effects of sex chromosome dosage on sex-chromosomal gene expression, we next investigated genome-wide responses to sex chromosome dosage. Using a linear model to estimate the slope of regression (β_X_ or β_Y_, *i.e.*, the log_2_ fold-change in expression per additional Xi or Y), we modeled log_2_ normalized read counts, controlling for sequencing batch. In CD4+ T cells, 202 autosomal genes responded significantly to Xi dosage, and 122 autosomal genes responded significantly to Chr Y dosage, with 56 genes responding significantly to both Xi and Chr Y dosage (268 unique Xi- and/or Y-responsive genes in all) (**Figure 3A-D; Table S4**). In monocytes, 54 and 46 autosomal genes responded significantly to Xi or Y dosage, respectively, with 11 genes responding significantly to both Xi and Y dosage (89 unique Xi- and/or Y-responsive genes in all) (**Figure 3E-H; Table S4**). For both primary cell datasets, saturation analyses revealed that there are likely more autosomal genes that respond to sex chromosome dosage to be found, should the dataset become better powered with samples from more individuals, as is the case for the LCL and fibroblast datasets (**Figure S4C-F**). Specifically, power analyses indicated that the current models detect approximately half of the significantly Xi-responsive autosomal genes in CD4+ T cells and two-thirds of those in monocytes (**Figure S7**).

**Figure 3.**
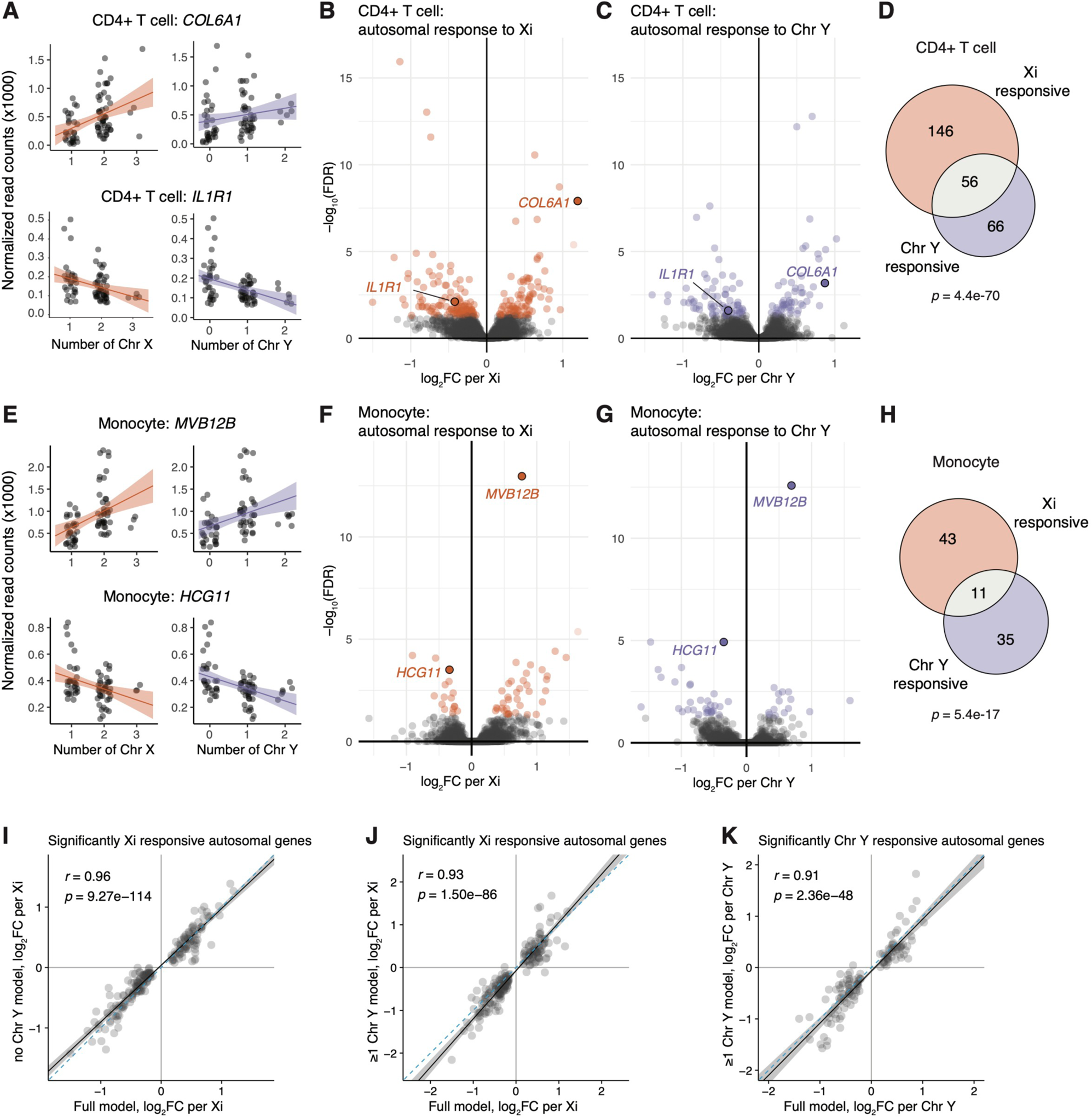
CD4+ T cells and monocytes show significant autosomal gene responses to Xi and Chr Y dosage. **(A)** Examples of genes positively and negatively responsive to Xi (left) or Chr Y (right) dosage in CD4+ T cells. Regression lines with confidence intervals are shown. **(B, C)** Volcano plots of autosomal responses in CD4+ T cells to Xi dosage (84 positively and 118 negatively responsive genes) **(B)** and Chr Y dosage (52 positively and 70 negatively responsive genes) **(C)**. **(D)** Overlap of autosomal genes with significant responses to Xi and Chr Y in CD4+ T cells; *p*-value from hypergeometric test. **(E)** Examples of genes positively and negatively responsive to Xi (left) or Chr Y (right) dosage in monocytes. **(F, G)** Volcano plots of autosomal responses in monocytes to Xi dosage (38 positively and 16 negatively responsive genes) **(F)** and Chr Y dosage (15 positively responsive and 31 negatively responsive genes) **(G)**. **(H)** Overlap of autosomal genes with significant responses to Xi and Chr Y dosage in monocytes; *p*-value from hypergeometric test. **(I-K)** In CD4+ T cells, correlations of log_2_ fold-changes per Xi (using full model: all six karyotypes) against Xi or Chr Y dosage in subset models as indicated. Solid black line indicates Deming regression slope, with 95% confidence intervals shaded gray. Pearson’s correlation coefficients shown; dashed blue lines indicate X=Y identity.

### Autosomal responses to sex chromosome dosage are robust to age and hormonal environment

To assess how much donor age impacts sex chromosome responsiveness in autosomal genes, we again generated an alternate linear model that included a term for age in addition to Chr X count, Chr Y count, and sequencing batch. This alternate model revealed 30 age-responsive genes in CD4+ T cells and five age-responsive genes in monocytes, but only two genes in CD4+ T cells (and no genes in monocytes) responded significantly to both age and sex chromosome dosage (**Figure S8, Table S5**). Including age in the linear model had little impact on the effect sizes of autosomal gene responses to Xi or Chr Y dosage (**Figure S8**). Accordingly, we modeled autosomal gene expression with no age term in our equations, just as we had done in modeling sex-chromosomal gene expression.

In human embryos, the presence of Chr Y leads to development of testes, which secrete testosterone, while the absence of Chr Y leads to development of ovaries, which secrete estrogen. To examine possible hormonal influences on the sex chromosome response, we re-assessed effects of Xi in samples that had no Chr Y (45,X; 46,XX; and 47,XXX) or that carried one Chr Y (46,XY and 47,XXY), as well as effects of Xi or Chr Y dosage in samples that had one or two Chr Ys (46,XY; 47,XXY; and 47,XYY). These subset models thus focus on samples with comparable hormonal profiles rather than a mix of estrogen and testosterone-dominant profiles. Despite reduced sample sizes in the subset models, autosomal responses to Xi and Chr Y dosage in the subset and full models were significantly correlated (**Figure 3I-K**, **Figure S9**). The similar responses across the full and subset models indicate that the autosomal response to sex chromosome dosage is not an artifact of the presence or absence of Chr Y; that is, the responses to sex chromosome dosage cannot be explained by estrogen versus testosterone alone.

### ZFX and ZFY drive a portion of the autosomal response *in vivo*

We showed previously that, in cultured cells, the impact of Xi on autosomal gene expression is similar to that of Chr Y.^12^ This suggested that the autosomal expression patterns were driven by shared factors on Chr X and Chr Y, *i.e.*, PAR genes or NPX-NPY gene pairs. We asked whether these *in vitro* findings and insights also held in primary CD4+ T cells and monocytes taken directly from the body. Indeed, autosomal gene responses to Xi and Chr Y dosage, in both CD4+ T cells and monocytes, were remarkably correlated for significantly responsive genes and across all expressed genes (**Figure 4A, B, S10A, B**). Thus, PAR genes and NPX-NPY homologous pairs are candidate drivers of the autosomal response both *in vitro* and *in vivo*.

**Figure 4.**
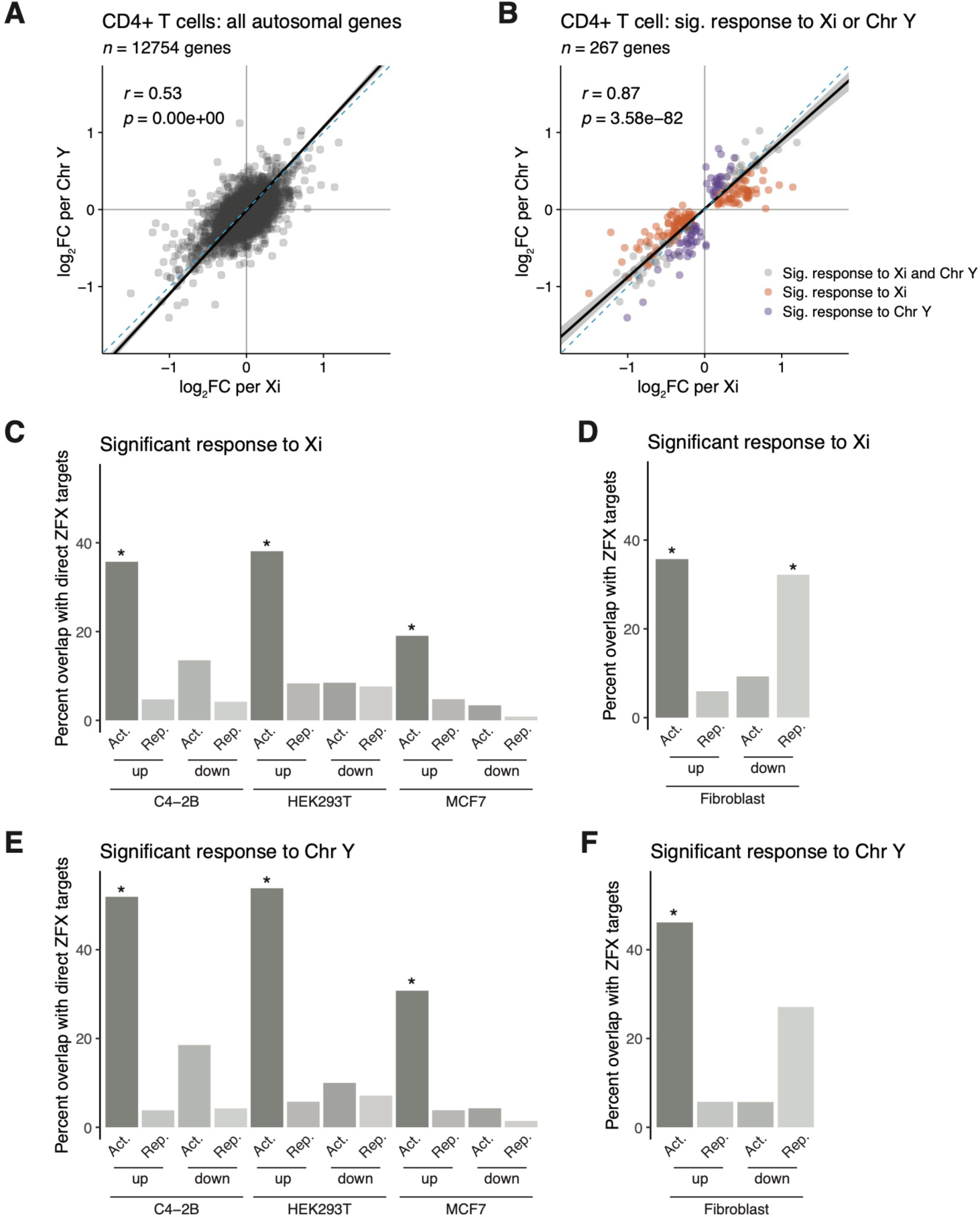
The transcription factor ZFX drives a significant fraction of autosomal response to sex chromosome dosage in immune cells. **(A, B)** Scatter plot of log_2_ fold-changes per Xi versus per Chr Y of all expressed autosomal genes in CD4+ T cells **(A)** or limited to genes significantly responsive to Xi dosage (orange), to Chr Y dosage (purple), or both (gray) **(B)**. **(C - F)** Bar plots showing percentages of genes with significant responses to Xi and/or Chr Y dosage in CD4+ T cells that were also identified as ZFX direct target genes in MCF7, C4-2B, and HEK-293T cell lines, or ZFX responsive in fibroblasts. Genes are parsed by whether they significantly increased or decreased with Xi or Chr Y dosage (“up” or “down”) and were activated or repressed by ZFX (“Act.” or “Rep.”). *p*-values reflect hypergeometric tests to identify significant enrichments of ZFX target genes in Xi- or Chr Y-responsive genes. Each comparison was restricted to genes expressed in both CD4+ T cells and the given *in vitro* cell line. Asterisks indicate the enrichments’ *p*-values were lower than the Bonferroni-adjusted threshold of 0.001.

Experiments in fibroblasts previously demonstrated a role for the transcription factors ZFX and ZFY, one of the ancestral NPX-NPY pairs, in driving much of the autosomal response *in vitro*.^12^ We set out to test whether changing the *ZFX* and/or *ZFY* expression level could also explain autosomal gene responses to Xi and Chr Y dosage *in vivo*. We first compared gene expression profiles from previously generated *ZFX* and/or *ZFY* CRISPRi knockdowns in fibroblasts to averaged autosomal responses to Xi and Chr Y dosage in CD4+ T cells and monocytes. Despite the distinct transcriptional profiles of fibroblasts versus CD4+ T cells and monocytes, ∼59% of the autosomal responders to Xi or Chr Y in CD4+ T cells and 56% of those in monocytes were significantly differentially expressed with *ZFX* and/or *ZFY* knockdown (*p* = 1.56e-06 and *p* = 0.01, respectively) (**Figure S10C, D**).

ZFX is a transcriptional activator that targets thousands of genes across the genome.^37,38^ We hypothesized that ZFX activates autosomal genes whose expression increases with Xi or Chr Y dosage. Utilizing lists of ZFX target genes (as determined by ZFX knockdown and ZFX binding) in C4-2B, HEK293T, and MCF7 cells, we sorted the targets into those whose expression decreases with ZFX loss (“ZFX activated”) or increases with ZFX loss (“ZFX repressed”).^37,38^ Autosomal genes whose expression increased with either Xi or Chr Y dosage in CD4+ T cells were significantly enriched for direct activation by ZFX (**Figure 4C, E**). In contrast, genes whose expression decreased with Xi or Chr Y showed no such enrichment (**Figure 4C, E**). When comparing Xi- or Chr Y-responsive genes to the CRISPRi ZFX knockdown in fibroblasts (which, lacking ZFX binding data, should represent a combination of direct and indirect targets of ZFX), we found significant enrichments for both i) ZFX-activated genes among the positively responsive genes and ii) ZFX-repressed genes among the negatively responsive genes (**Figure 4D, F**). While we observed similar trends among genes that responded positively to Xi or Y dosage in monocytes, these trends were not statistically significant following multiple hypothesis corrections (**Figure S10E-H**). These findings indicate that ZFX and ZFY drive a significant portion of the autosomal response to sex chromosome dosage both *in vitro* and *in vivo* – and that additional sex-chromosomal drivers of autosomal gene expression remain to be elucidated.

### Autosomal responders to Xi and Chr Y dosage show cell-type-specific functional enrichments

We next examined the annotated functions of the autosomal genes most responsive to Xi and Chr Y dosage in CD4+ T cells and monocytes. Gene set enrichment analysis of the Hallmark pathways, as defined by the Human Molecular Signatures Database, revealed several significantly enriched gene sets in each primary immune cell type (**Figure 5A; Table S6, S7**).^39,40^ CD4+ T cells and monocytes had largely distinct pathway enrichments. In CD4+ T cells, interferon alpha (IFNα) and gamma (IFNγ) response genes, complement, and metabolic pathways such as fatty acid metabolism and oxidative phosphorylation all had significantly negative responses to Xi and Chr Y dosage; *i.e.*, expression of these gene sets decreased with increasing numbers of X or Y chromosomes (**Figure 5A**). In contrast, monocytes showed significant positive enrichments for pathways such as “TNFα signaling via NF-κB” and “inflammatory response” (**Figure 5A**). Notably, significant enrichments for IFNα and IFNγ responses in CD4+ T cells persisted across subset models restricted to karyotypes with or without a Chr Y, indicating that the negative response to sex chromosome dosage is not driven merely by the presence or absence of a Chr Y (or, by proxy, a testosterone-dominant hormonal milieu) (**Figure S11; Table S6**).

**Figure 5.**
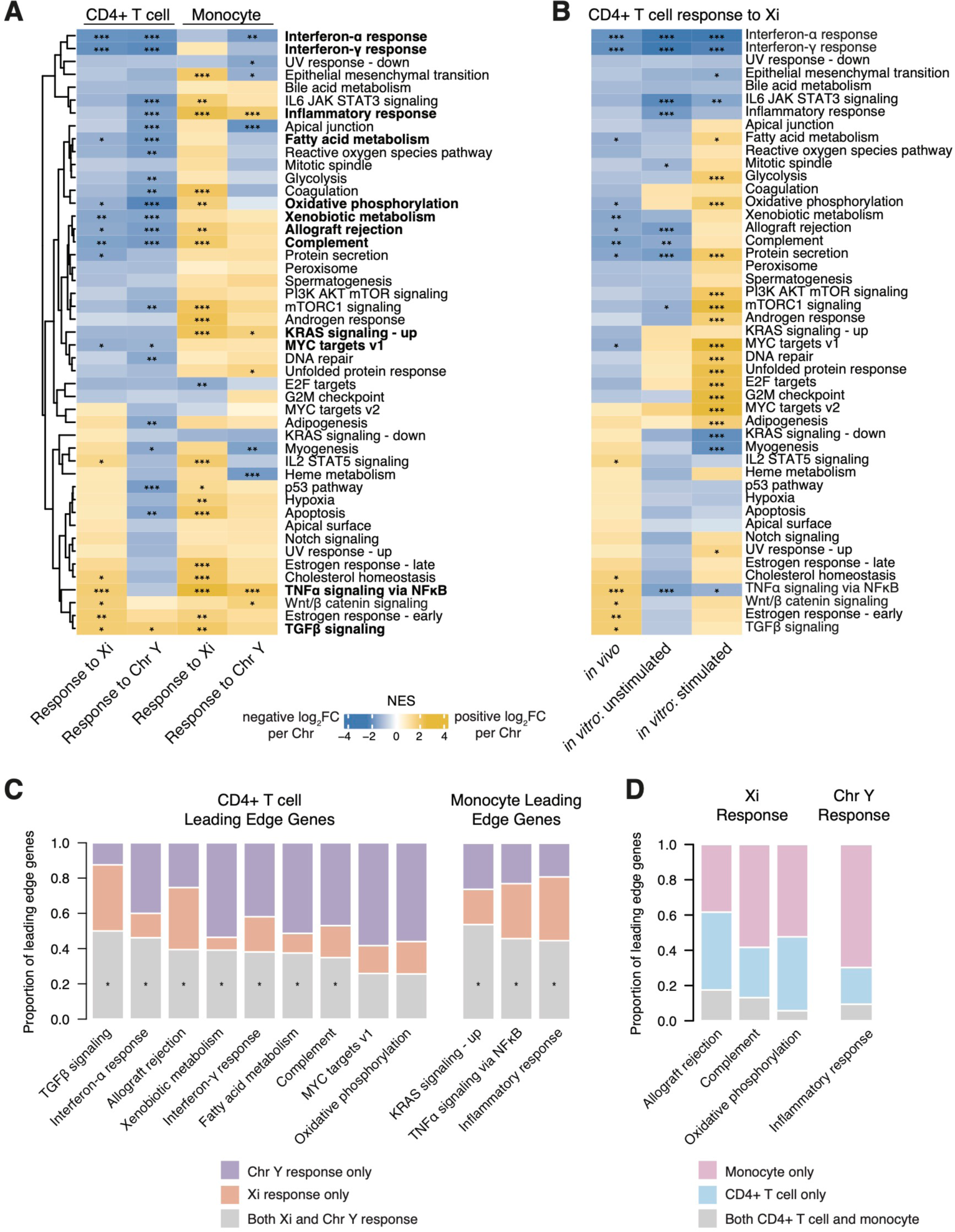
Autosomal responders to Xi and Chr Y dosage are functionally enriched for interferon and TNFα signaling pathways. **(A)** Heatmap of normalized enrichment scores (NES) for Hallmark gene sets in CD4+ T cells and monocytes; significant enrichments indicated by asterisks (* *p*-adj <0.05; ** *p*-adj < 0.01; *** *p*-adj < 0.001). CD4+ T cell pathways and monocyte pathways with significantly concordant Xi and Chr Y responses are in bold. **(B)** Heatmap of normalized enrichment scores for Hallmark gene sets in CD4+ T cells *in vivo* and unstimulated and stimulated CD4+ T cells *in vitro*; significant enrichments indicated by asterisks, as above. **(C)** Proportions of leading edge genes, in the indicated pathways, that are specific to the Xi response (orange), specific to the Chr Y response (purple), or common to Xi and Chr Y responses (gray). Asterisks indicate significant overlaps of leading edge genes between Xi and Chr Y responses. **(D)** Proportions of leading edge genes, in indicated pathways, that are specific to CD4+ T cells (blue), to monocytes (pink), or shared by the two cell types (gray).

To assess the effects of Xi dosage on cell function, we activated CD4+ T cells from individuals with one or two X chromosomes (45,X; 46XY; 46,XX; and 47,XXY karyotypes) *in vitro* with beads biotinylated with CD2, CD3, and CD28 antibodies and then performed RNA-Seq (**Figure S12; Table S8, S9**). Many transcriptional patterns persisted between the *in vivo* and unstimulated *in vitro* CD4+ T cells, including decreased expression of IFNα and IFNγ response genes with increasing Xi copy number (**Figure 5B, Table S10**). Upon activation, we observed even starker effects of Xi dosage on gene expression, with several cell-cycle-associated gene sets, such as MYC targets, E2F targets, and G2M checkpoint genes, significantly increased with Xi dosage. These results demonstrate functional consequences of Xi dosage in immune cells and suggest that further sex chromosomal effects may be revealed in other states of cell stimulation or perturbation.

We next asked whether Xi dosage and Chr Y dosage similarly impacted the enriched pathways within a given cell type. In pathways with significant responses to both Xi and Chr Y, the genes responsible for these results were largely the same. There were significant overlaps between the sets of “leading edge” genes responsive to Xi and to Chr Y, with generally correlated effect sizes (**Figure 5C, Table S11**). While these results underscore the broadly similar effects of Xi and Chr Y dosage on autosomal genes within cell types (**Figure 4A, B, Figure S10A, B**), regressions between Xi and Chr Y effects for several pathways were skewed, with unequal responses to Xi versus Chr Y dosage (*i.e.*, pathways with slopes ≠ 1) (**Table S11**).

In addition to gene pathways that were significantly enriched in only one of the two immune cell types, we observed pathways in which sex chromosome dosage had opposite effects in different cell types. These included the complement and “inflammatory response” gene sets, whose expression increased with sex chromosome dosage in monocytes and decreased in CD4+ T cells (**Figure 5A**). The “leading edge” genes responsible for the significant enrichments in CD4+ T cells versus monocytes for these gene sets were largely cell-type-specific, with no significant overlap between the two cell types (**Figure 5D)**. This indicates that, rather than having opposing effects on the same genes across cell types, sex chromosome dosage impacts different genes within these pathways across cell types. Taken together, these results underscore the cell-type specificity of responses to Xi and Chr Y dosage.

### Autosomal responses are more cell-type-specific than are sex-chromosomal responses

To investigate differences and similarities between cell types in autosomal responses to Xi and Y dosage, we first examined genes with statistically significant responses. Out of 247 autosomal genes that responded to Xi dosage in CD4+ T cells and/or monocytes, only nine were significantly responsive in both immune cell types; of 1,857 autosomal genes that responded to Xi dosage in one or more of the four cell types, only two were significantly responsive across CD4+ T cells, monocytes, LCLs, and fibroblasts (**Figure 6A**). X-chromosomal responses to Xi dosage were decidedly more consistent: of 124 X-chromosomal (including PAR and NPX) genes that had a significant ΔE_X_ in at least one of the four cell types, 19 genes responded significantly to Xi dosage in all four cell types (**Figure 2D**). Even more strikingly, five of the seven NPY genes that were expressed in all four cell types had a significant ΔE_Y_ in all four cell types, but only one autosomal gene – out of 652 – had a significant response to Chr Y dosage in all four cell types (**Figure S6, Figure 6B**).

**Figure 6.**
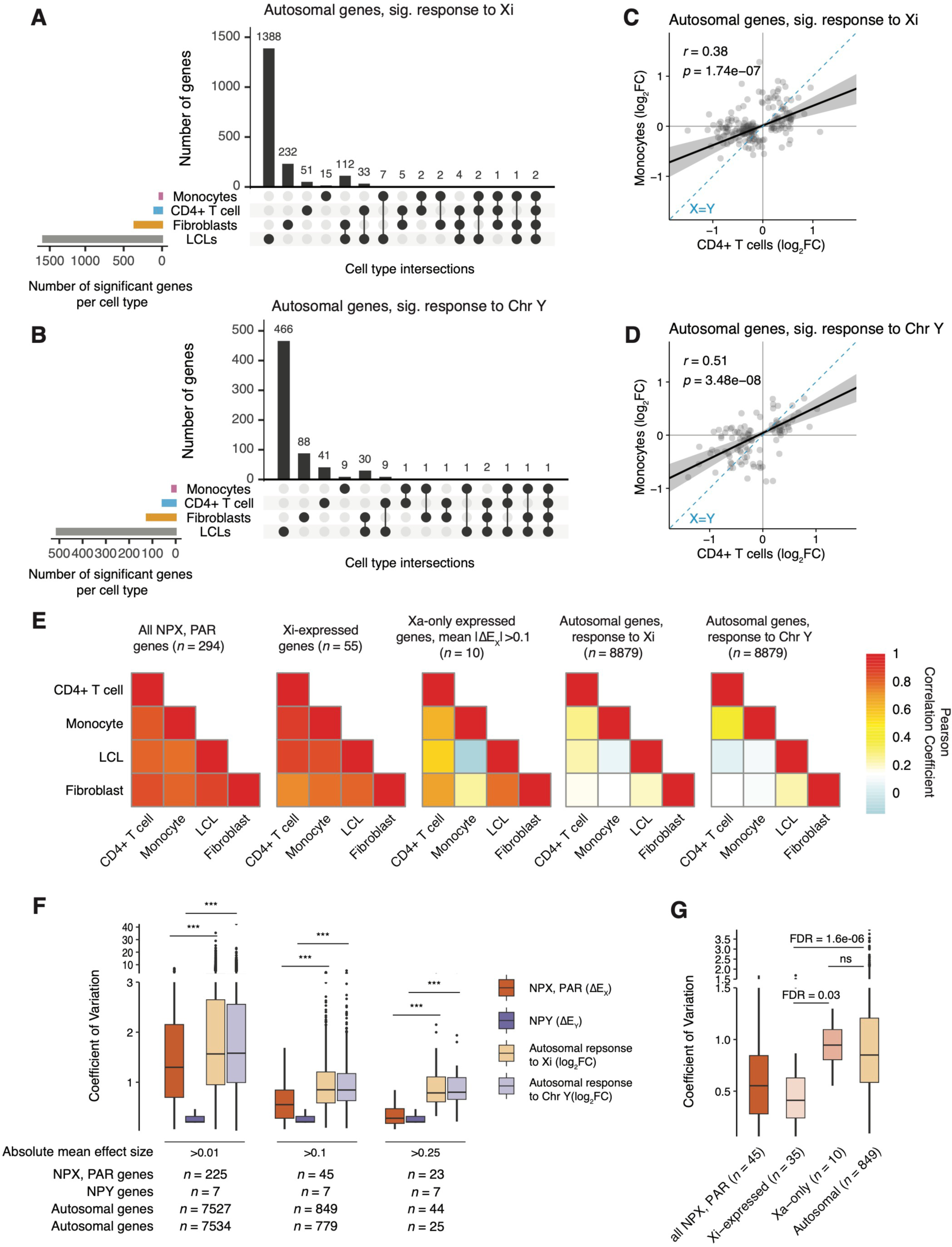
Autosomal responses to Xi and Chr Y dosage are more cell-type-specific than are sex-chromosomal responses. **(A, B)** UpSet plots showing limited commonalities across cell types of autosomal genes with significant responses to Xi **(A)** or Chr Y **(B)**. Analyses restricted to genes expressed in all four cell types. **(C, D)** Correlations of Xi **(C)** and Chr Y **(D)** log_2_ fold-changes between CD4+ T cells and monocytes. Plotted genes reached statistical significance in either CD4+ T cells or monocytes. Deming regression indicated by solid black line with 95% confidence intervals shaded gray; dotted blue line indicates X=Y identity. Pearson correlation coefficients shown. **(E)** Heatmaps of Pearson correlation coefficients between ΔE_X_ values of all NPX and PAR genes across four cell types (left); between ΔE_X_ values of all Xi-expressed genes across four cell types (second from left); between ΔE_X_ values of Xa-only expressed genes, with mean absolute ΔE_X_ values of at least 0.1, across four cell types (middle); and between log_2_ fold-changes of all expressed autosomal genes in response to Xi dosage (second from right) or Chr Y dosage (right). Analyses restricted to genes expressed in all four cell types. **(F)** Variation in gene-by-gene responses to Xi or Y dosage, as indicated, across cell types was calculated by finding coefficients of variation (CV = standard deviation in effect size across cell types / absolute mean effect size across cell types) for genes expressed in all four cell types. Box plots show median (line), interquartile range (IQR; top and bottom of box), and 1.5 * IQR (whiskers) of CV values. Gene sets limited to those with absolute mean effect sizes above the indicated values; FDR calculated from unpaired t-tests and asterisks indicate FDR < 1e-06. **(G)** Coefficients of variation for responses to Xi dosage for genes in the indicated classes. Analyses restricted to genes expressed in all four cell types with an absolute mean effect size >0.1. Box plots show median (line), interquartile range (IQR; top and bottom of box), and 1.5 * IQR (whiskers) of CV values. FDR calculated from unpaired t-tests.

Although only nine autosomal genes responded significantly to Xi dosage (and only six genes to Chr Y dosage) in both CD4+ T cells and monocytes, the magnitude and direction of response to Xi (or Chr Y) dosage were correlated across all 11,142 expressed autosomal genes – and these correlations became stronger when limited to significant autosomal responders (**Figures 6C, D, S13**). However, even among significantly Xi-responsive genes, the correlation between CD4+ T cell and monocyte autosomal gene responses was weaker than X-chromosomal ΔE_X_ correlations between the same cell types. This pattern persisted across all pair-wise cell type comparisons, with ΔE_X_ values of X-chromosomal genes highly correlated between cell types, whereas autosomal gene responses were less concordant (Pearson correlation coefficients ranging from 0.07 to 0.27; **Figure 6C-E, S13**). The strong ΔE_X_ correlations between NPX and PAR genes persisted when restricting these analyses to genes expressed on Xi, while genes only expressed on Xa had more cell-type-dependent responses to Xi dosage (**Figure 6E**). Thus, while X- and Y-chromosomal responses to Xi and Chr Y dosage are preserved across cell types, downstream effects of Xi and Chr Y dosage on autosomal and Xa expression are more cell-type-specific. This comports with recent reports on cultured cells, where Chr X’s effects in *cis* appeared more stable than those in *trans*.^11,12,41^ Specifically, we find that the responses of autosomal genes to Xi dosage are as much as 2.6 times as variable (between cell types) as those of NPX and PAR genes (FDR = 9.07e-09; mean coefficient of variation [CV] for autosomal genes with mean absolute effect sizes > 0.25 is 0.90; CV for NPX and PAR genes is 0.35) (**Figure 6F**). Parsing X-chromosomal genes by Xi expression, we found that Xi-expressed genes’ variation in response to Xi dosage is significantly lower than that of genes not expressed on Xi (“Xa-only”), while the variability in responses of Xa-only genes was not statistically different from that of autosomal genes (FDR = 0.97). (**Figure 6G**). Finally, NPY gene responses to Chr Y dosage showed lower variation than autosomal responses to Chr Y dosage (FDR = 0.001) (**Figure 6F**). In summary, Xi-expressed genes and NPY genes show little variation in this regard across cell types, while Xa-only genes display variation as high as that seen with autosomal genes.

## Discussion

We explored the transcriptome-wide impact of Xi and Chr Y dosage *in vivo*, employing linear modeling of gene expression in primary immune cells of individuals with sex chromosome aneuploidy. We characterized Xi- and Chr Y-responsive genes *in vivo* and quantified similarities and differences in these responses among four human cell types. For sex-chromosomal gene expression, the metrics ΔE_X_ and ΔE_Y_ quantify the impact of Xi and Chr Y with gene-by-gene specificity. The wide range of ΔE_X_ values across X-chromosomal genes – within a single cell type – underscores the importance of such a metric to gauge Xi’s contributions to X-chromosomal expression on a gene-by-gene basis. By contrast, the relative consistency of ΔE_Y_ values across expressed Y-chromosomal genes speaks to the absence of special regulatory mechanisms on supernumerary Y chromosomes. Linear modeling also revealed abundant *trans* effects on autosomal gene expression.

### The influence of Xa, Xi, and Y on human immune cell function

Given the variety and extent of sex differences in immune function, immune cells present an especially poignant opportunity to dissect the extent and nature of Xa, Xi, and Y chromosomal influence on human gene expression and cell function. Evidence indicates that both hormones and sex chromosomes drive sex-biased phenotypes, but their roles have not been fully disentangled.^42–44^ Previous studies of sex chromosomal influences on immune cell function have revealed immune dysfunction in sex chromosome aneuploid individuals – including increased risk of autoimmunity in Turner Syndrome (45,X) and Klinefelter Syndrome (47,XXY) – while other molecular studies have focused on the effects of X-linked immune genes that are expressed from both Xa and Xi.^45–48^ For example, the X-linked toll-like receptor 7 gene, *TLR7*, is biallelically expressed in some human 46,XX plasmacytoid dendritic cells, B cells, and monocytes, and in turn has been associated with sex-biased IFNα and IFNγ responses and female-biased autoimmunity.^28,47–49^ Similarly, it has been proposed that heterogenous expression of other X-linked genes from Xi, including *CD40LG,* contributes to the female-biased prevalence of lupus.^31^ In our present study, we did not observe a significant increase in *TLR7* transcript levels in monocytes, nor increased *CD40LG* transcript levels in CD4+ T cells, with additional copies of Xi. These findings run counter to prior expectations that expression from Xi would result in increased total expression with increased Xi dosage. In our bulk RNA-Seq analyses, expression of *TLR7* and *CD40LG* from Xi is too modest to have a discernible effect on total expression levels.

In addition to studying the transcriptional profiles of CD4+ T cells *in vivo*, we measured the effects of Xi dosage on transcription upon CD4+ T cell activation *in vitro*. We discovered effects on gene expression that persisted *in vivo* and *in vitro,* including the IFNα and IFNγ response pathways, as well as context-specific gene expression patterns, such as TNFα and IL6 JAK/STAT3 signaling. Activation of CD4+ T cells *in vitro* revealed a host of additional functions impacted by Xi dosage. Upon stimulation, CD4+ T cells that possessed an Xi had higher expression of proliferative and metabolic genes than CD4+ T cells lacking an Xi. These results highlight specific pathways through which Xi may influence CD4+ T cell responses to activation and, in turn, contribute to sex-biased immune phenotypes.

### Xi, Xa, and Y: an emerging understanding of gene regulation in cis and trans

Having systematically analyzed the expression of protein-coding Xa, Xi, Y, and autosomal genes across four cell types, we can identify some general patterns regarding the sex chromosomes’ wide-ranging roles in regulating the genome. Four major findings appear to hold *in vivo* and *in vitro*, and they generalize across cell types: 1) Xi-expressed genes display stable ΔE_X_ values, 2) Y-chromosomal genes display stable ΔE_Y_ values, 3) Xa and autosomal responses to Xi and Y display cell-type specificity, and 4) Xa and autosomal responses are driven by a small number of widely expressed NPX-NPY gene pairs.

First, we observe a striking consistency in the responses of Xi-expressed genes to Xi dosage, such that, gene by gene, ΔE_X_ values are remarkably stable across cell types, both *in vivo* and *in vitro*. The simplest interpretation is that, for each Xi-expressed gene, the ratio of Xi to Xa transcription is conserved among somatic cell types. The molecular determinants of these ratios and their stability are presently unknown.

Second, we see a comparable or even greater consistency in the responses of Y-chromosomal genes to Chr Y dosage, with ΔE_Y_ values for each gene being strikingly similar across cell types, *in vivo* and *in vitro*. With median ΔE_Y_ values of broadly expressed NPY genes ranging from 0.65 to 1.07, our findings are consistent with the absence of Y inactivation or other dosage compensation mechanisms *in vivo* and *in vitro*.

Third, while transcription from Xi and Chr Y is stable and robust across cell types, the transcriptional responses of autosomal and Xa-only genes are cell-type-specific. In this critical epigenetic respect, Xa behaves like an autosome and not like its genetically identical twin, Xi. Remarkably, the Xi and Y chromosomes can elicit cell-type-specific responses on autosomes and Xa even when the Xi and Y transcriptomes do not vary substantially among cell types.

Fourth, evidence is growing that a small, fixed set of Xi- and Y-expressed genes drive these cell-type-specific autosomal and Xa responses to Xi and Chr Y dosage. In each of the four studied cell types, Xi and Chr Y have comparable effects on genome-wide expression, indicating that regardless of cellular identity, factors expressed by both Xi and Chr Y drive most of the genome-wide response. The observation that Xi vs. Chr Y effects are *similar* but not *identical* (**Figure 4A, B, Figure S10A, B**)^12^ aligns well with evidence ^12^implicating the diverged NPX-NPY genes, such as *ZFX* and *ZFY*, rather than PAR genes, which are identical on Chrs X and Y. We hypothesize that the cell-type-specificity of autosomal and Xa responses to conserved Xi- and Y-expressed drivers is the result of differences between cell types in chromatin accessibility and autosomal cofactors. This hypothesis should now be explored experimentally.

Taken together, these findings provide a counterpoint and complement to the well-established concept of “variable escape” from X chromosome inactivation, which posits that the set of X-chromosomal genes expressed from Xi varies substantially across cell types.^5,33,36,50,51^ While our study of four cell types does not exclude the occurrence of biologically critical variation in the Xi transcriptome among these and other cell types, our findings do not provide direct evidence of such variation. Instead, our findings show that the response of autosomal and Xa-only genes is the more cell-type-specific phenomenon – at least among the four cell types that we have analyzed. Studies of additional somatic cell types will be required to resolve this important question.

## Conclusion

Through linear modeling of a diverse array of naturally occurring sex chromosome constitutions, we have characterized quantitatively the *in vivo* impact of Xi and Chr Y on human gene expression, in two immune cell types. We find consistent, conserved regulation of Xi and Chr Y gene expression across distinct cell types but divergent effects of Xi and Chr Y on autosomal and Xa gene expression. Xi and Chr Y have comparable regulatory effects on transcription genome-wide, both *in vitro* and *in vivo*, driven in part by the X- and Y-encoded homologous pair of transcription factors *ZFX* and *ZFY*. The studies reported here establish a foundation for fully dissecting how Xi and Chr Y affect transcription and cell function throughout the body.

## Acknowledgements

We thank members of the Page Laboratory, especially Daniel W. Bellott, for critical reading and comments on the manuscript, and Jorge Adarme and Susan Tocio for laboratory support. We thank the Whitehead Genome Technology Core facility for library preparation and sequencing.

## Funding

Lallage Feazel Wall Damon Runyon Cancer Research Foundation Fellowship (LVB)

National Institutes of Health grant F32HD091966 (AKSR)

Simons Foundation Autism Research Initiative – Collaboration Award (DCP, JFH)

National Institutes of Health grant U01HG0007587 (DCP and MM)

Howard Hughes Medical Institute (DCP)

National Human Genome Research Institute Intramural Research Program (MM, DK)

Contents are the authors’ sole responsibility and do not necessarily represent official NIH views.

Philanthropic support from:

The Brit Jepson d’Arbeloff Center on Women’s Health

Arthur W. and Carol Tobin Brill

Matthew Brill

Charles Ellis

The Brett Barakett Foundation

Howard P. Colhoun Family Foundation

Seedlings Foundation

Knobloch Family Foundation

## Author Contributions

Conceptualization: LVB and DCP

Data curation: LVB and AKSR

Formal analysis: LVB, AKSR, and HS

Funding acquisition: LVB, AKSR, JFH, MM, and DCP

Investigation: LVB, GW, TTP

Methodology: LVB, GW, AKSR, HS

Project administration: LVB, GW, JFH, LB, AKSR, AB, PK, and NB

Resources: AB, PK, NB, AEL, and MM

Software: LVB, AKSR, AKG, and HS

Supervision: LVB, AKSR, JFH, LB, AEL, DK, MM, and DCP

Validation: LVB, AKSR, and HS

Visualization: LVB

Writing - original draft preparation: LVB and DCP

Writing - review and editing: LVB, AKSR, JFH, and DCP

## Inclusion and diversity

We support inclusive, diverse, and equitable conduct of research.

## STAR Methods

### RESOURCE AVAILABILITY

#### Lead contact

Further information and requests for resources and reagents should be directed to and will be fulfilled by the lead contact, David C. Page.

#### Materials availability

There are restrictions on the availability of the CD4+ T cell and monocyte cells in this study, as the studied cells are primary cells that have not been stably transformed into cell lines.

#### Data and code availability

- Raw RNA-Seq data has been deposited to dpGAP and processed data has been deposited at github. Both are publicly available as of the date of publication. Accession numbers and DOIs are listed in the key resources table.
- This paper analyzes existing, publicly available data. Accession numbers for these datasets are listed in the key resources table.
- Original code has been deposited at github and is publicly available as of the date of publication. The accession number is listed in the key resources table.
- Any additional information required to reanalyze the data reported in this paper is available from the lead contact upon request.

### EXPERIMENTAL MODEL AND STUDY PARTICIPANT DETAILS

#### Human subjects

Recruitment of euploid and sex chromosome aneuploid individuals has been previously described, but is restated briefly here: Seventy-two adults (18+) were recruited through a joint IRB-approved study between the NIH Clinical Center (12-HG-0181) and Whitehead Institute; 3 adults were recruited through Massachusetts General Hospital and 4 adults were recruited through Boston Children’s Hospital (Protocol #1706013503). Informed consent was obtained from all participants. Extensive karyotyping was performed on primary blood cells derived from sex chromosome aneuploid individuals recruited through the NIH clinical center; of 1000 karyotyped cells, the dominant karyotype for the diagnosed aneuploidy was, on average, 94% ± 8.7% of cells.

#### Isolation of Peripheral Blood Mononuclear Cells

Whole blood was collected in BD Vacutainer Sodium Heparin tubes (GTIN: 00382903678747) and enriched for CD4+ T cells and monocytes on the same day as collection. First, peripheral blood mononuclear cells were isolated by transferring whole blood to a 50 mL conical tube and adding 1X PBS without calcium and magnesium to a total volume of 40 mL. After capping the tube and inverting the blood mixture 5 – 6 times, 10.5 mL of lymphocyte separation media (LSM; MP Biomedicals cat. #50494) was added to the to the tube by placing the pipette directly into the blood mixture and slowly releasing the LSM by gravity. The blood mixture was spun at 1500 rpm for 30 minutes, at room temperature, with no brake. Without disturbing the separated layers, the cloudy buffy coat of PBMCs were transferred to a new tube and washed with 1X PBS for 10 minutes at 1500 rpm at room temperature.

#### Enrichment of CD4+ T cells and monocytes from PBMCs

PBMC cells were split in equal volumes and enriched for CD4+ T cells using the Miltenyi Biotec Human CD4+ T Cell Isolation Kit (cat# 130-096-533) and for monocytes using the Miltenyi Biotec Human Monocyte Isolation Kit (cat# 130-091-153), following the manufacturer’s protocol. Following enrichment, each cell type was resuspended in freezing media (FBS + 10% DMSO) and stored at −80C° until RNA extraction.

### METHOD DETAILS

#### RNA extraction

For RNA extraction, 1 million cells per sample were spun down for 5 minutes at 13,000rpm and resuspended in 0.5 mL TRIzol Reagent (Life Technologies #15596026). Following a 5 minute incubation at room temperature, each sample was transferred to a 2 mL 5Prime Gel Heavy PhaseLock tube (Quantabio #2302830) and given 0.1 mL of chloroform. Samples were then incubated for 5 minutes at room temperature on a Thermomixer (Eppendorf) at 1100rpm, then spun for 5 minutes at 13,000rpm, room temperature. The aqueous phase, containing RNA, was transferred to a new PhaseLock tube and given 0.2 mL acid phenol:chloroform (Life Technologies AM 9722), followed by a second 5 minute Thermomixer incubation. Samples were again spun to separate phases, and the aqueous phase was again transferred to a new Eppendorf tube. RNA was precipitated overnight at −20C with 0.5 uL Glycoblue (Life Technologies AM9515), 20 uL 3M sodium acetate, pH 5.5, and 0.5 mL RNase-free ethanol, and resuspended in 15 uL RNase-free water. RNA concentrations were quantified using the RNA HS Qubit kit (ThermoFisher).

#### RNA sequencing and analysis

RNA sequencing libraries were prepared at the Whitehead Genome Technology Center, using the KAPA HyperPrep kit, followed by size selection for 300-600bp fragments with the PippinHT system (Sage Science) and 2% agarose gel. Paired 100×100bp reads were sequenced on the Illumina HiSeq 2500 or NovaSeq 6000. Following RNA sequencing, data was pseudoaligned using kallisto (v0.42.5), using the --bias flag, to a custom human genome based on the GENCODE v24 Basic annotation, as detailed previously.^11,12^ kallisto results were imported into R for downstream analysis using the tximport package. We restricted our analysis to protein-coding genes and lincRNAs, with the exception of also including the Y-chromosomal annotated pseudogenes *PRKY* and *TXLNGY*. We filtered for expression by a median TPM >1 in either XX or XY samples within a given cell type.

#### CIBERSORTx analysis of cell type populations

To validate successful CD4+ T cell and monocyte enrichment, CIBERSORTx was used to impute cell fractions from the bulk RNA-Seq samples.^25^ Following kallisto pseudoalignment, TPMs were extracted using the tximport package and uploaded to CIBERSORTx (https://cibersortx.stanford.edu/). Cell fractions were imputed using the LM22 signature matrix file as reference for 22 immune cell types, with batch correction (B-mode) enabled.

#### Linear modeling of sex-chromosomal genes

Linear models for each expressed X-chromosomal and Y-chromosomal gene were generated using the lm() function in R, as previously described in San Roman, et al., 2023 and briefly restated here. For NPX and PAR genes, normalized read counts from all samples were inputted to the model, with terms for the number of Xi, the number of Chr Y, and sequencing batch. For NPY genes, karyotypes with at least 1 Chr Y were inputted to model the number of Xi, the number of Chr Y minus one, and sequencing batch.

ΔE_X_ and ΔE_Y_ values were calculated by dividing the β_X_ or β_Y_, respectively, by the corresponding β_0_ (*i.e.*, the intercept, representing expression from 45,X samples for NPX and PAR linear models and from 46,XY and 47,XXY samples for NPY linear models). To calculate ΔE_X_ and ΔE_Y_ for *XIST* (which is only expressed in karyotypes with at least two Chr X), β_X_ or β_Y_ were divided by β_0 +_ β_X_. P values were adjusted for multiple hypothesis correction using the Benjamini-Hochberg method; genes with *p*-adjusted < 0.05 were judged to be significant.

For gene classes, we defined PAR genes as those from PAR1. “NPX genes with NPY homolog” were defined as non-PAR X-chromosomal genes with an expressed NPY homolog (including NPY homologs currently annotated as pseudogenes). Note that for plotting purposes, the ΔE_X_ values for four X-chromosomal genes (*ZFX-AS1*, *PRKX-AS1*, and *EIF1AX-AS1* in CD4+ T cells and *BEX1* in fibroblasts) with outlying ΔE_X_ values greater than 2 – often inflated from low expression that led to small β_0_ denominators –were excluded from figures.

To examine the impact of age on the response to Xi or Chr Y, identical linear models were generated with the exception that age, in years, of the donor at time of sample collection was included as an additional term in the model.

#### Linear modeling of autosomal genes

For autosomal genes, log_2_ normalized read counts were used to model for Chr X number, Chr Y number, and sequencing batch via DESeq2; the “results” function within DESeq2 was then used to identify specific responses to Xi or Chr Y. Genes with adjusted *p*-values < 0.05 (as output by DESeq2) were called as significantly responsive to Xi or Chr Y. To examine the impact of age on the response to Xi or Chr Y, identical DESeq2 linear models were generated with the exception that age, in years, of the donor at time of sample collection was included as an additional term.

#### Saturation analysis

Separate saturation analyses were performed for X-chromosomal genes (NPX and PAR) and autosomal genes, consistent with the linear modeling approaches used to generate the sex-chromosomal versus autosomal models as described above. Size-*n* subsets were randomly generated, without replacement, 100 times for each sample size, *n*, and model matrixes were assessed to be full rank. Genes with adjusted *p*-value <0.05 were called as significant.

#### Power analysis

To estimate the power of the linear models to identify all significantly responsive genes, read counts for 100 genes from the sex chromosome aneuploidy cohort were simulated to generate effect sizes (log_2_ fold-changes) ranging from 0.05 to 1. To simulate the data, karyotype labels were first shuffled to remove expression dependencies on Chrs X or Y count, and then new, artificial read counts were generated such that for a given set of 100 random genes, read counts were altered to reflect the desired effect size. DESeq2 models on the simulated data were iteratively run 20 times for each effect size and compared with the actual DESeq2 results to find the fraction of true positive results (TPRs). After calculating median TPRs for each effect size in response to either Xi or Chr Y, in each cell type (CD4+ T cells and monocytes), power for each effect size was estimated by fitting an exponential curve. Using the curved fit, we then estimated the number of unobserved genes by applying the power for each effect size to the number of observed significantly responsive genes, by cell type and sex chromosome dosage.

#### Annotation of Xi-expressed and Xa-only expressed genes

A meta-analysis of previously published annotations of X-inactivation status was described in detail in San Roman, *et al.*, 2023 (summarized in Table S6 of that study) and further extended in San Roman, *et al.,* 2024.^11,12^ We used the expanded analysis in Table S4 from San Roman, *et al*., 2024 to annotate genes in the present study. Briefly, allelic ratio (AR) data was summarized across five studies utilizing various AR methodologies, including data from cDNA SNP-chips in skewed LCLs and fibroblasts, single-cell RNA-Seq in LCLs, single-cell RNA-Seq in fibroblasts, and allele-specific bulk RNA-Seq in LCLs and fibroblasts with skewed X-inactivation.^5,11,27,33–36^ Genes with evidence of expression from Xi in more than half of the studies, as well as those with AR ≥ 0.1 and with evidence of expression from Xi in at least one study, were designated as “Xi-expressed”. Genes that showed no expression from Xi across all studies, as well as those showing no Xi expression in at least half of studies, plus ARs < 0.1, were called as “Xa-only expressed”. In all, 107 genes are annotated as “Xi-expressed” and 460 genes as “Xa-only expressed”; 398 genes were not captured in the analyzed studies (“no call”).

#### ZFX direct target enrichment analysis

To generate lists of ZFX direct target genes, we utilized ZFX ChIP-Seq data from four cell lines available via ENCODE4 v1.6.1 (HCT 116, C4-2B, MCF-7, and HEK-293T), as well as RNA-Seq of ZFX knockdown experiments in the same cell types from ENCODE or GEO and RNA-Seq data from ZFX knockdown in fibroblasts.^12,37,38^ When available, we downloaded pre-processed RNA-Seq data, but otherwise processed fastq files using kallisto. To identify ZFX binding at promoters, we used the BEDTools closest function to map ZFX peaks to the nearest annotated transcription start site in the GENCODE v24 comprehensive annotation. Genes that 1) had ZFX bound to the promoter and 2) were significantly differentially expressed upon ZFX knockdown were classified as “direct target genes”. After direct ZFX target genes were identified for each cell type, other genes that decreased expression with ZFX loss were classified as “ZFX activated”, while genes that increased expression with ZFX loss were classified as “ZFX repressed”. We then found the intersection of “ZFX activated” and “ZFX repressed” genes with genes that either increase or decrease with Xi or Chr Y dosage, and performed hypergeometric tests using the phyper function in R to establish whether there was a significant overlap between gene lists.

#### Gene set enrichment analysis

Gene set enrichment analysis was conducted in R with the fgsea package, using the 50 Hallmark pathways (v7.1) downloaded from the Molecular Signatures Database (https://www.gsea-msigdb.org/gsea/msigdb).^39,40^ Analysis was restricted to autosomal protein-coding and lincRNA genes, which were ranked by each gene’s t-statistic from the DESeq2 models for Chr X or Y count.

#### In vitro activation of CD4+ T cells

Frozen CD4+ T cell stocks from individuals with 45,X, 46,XX, 46,XY, or 47,XXY karyotypes were thawed for 2-3 minutes in a 37°C water bath, followed by gentle addition of media (RPMI + 10% AB serum) to dilute the DMSO in the freezing media. Cell viability and concentrations were measured on a Countess Cell Counter. Cell concentrations were then normalized across samples to 2.77 × 10^6^ cells/mL, such that final culture concentrations (following the addition of activation beads or media) would be 2.5 × 10^6^ cells/mL. Activation beads were prepared following manufacturer’s instructions (Miltenyi T cell Activation/Expansion kit, human, cat.# 130-091-441). Cells were plated in a flat-bottom 96-well culture plate (∼300 uL/well) and either cultured in RPMI + 10% AB serum + IL-2 alone, or RPMI + 10% AB serum + IL-2 + biotinylated activation beads, following manufacturer’s instructions. After plating, cells were incubated at 37C° +5% CO_2_ for 24 hours prior to collection for RNA-Seq. RNA was extracted from each sample using the Zymo Quick RNA Microprep kit (cat. R1050), and RNA-Seq libraries were made via the IDT xGen RNA Library Preparation kit (cat. 10009814).

### QUANTIFICATION AND STATISTICAL ANALYSIS

We used various statistical tests to calculate *p*-values as indicated in Method Details, figure legends, or text, where appropriate. To calculate all statistics and generate plots, we used R software, version 3.6.3^31^. We considered results statistically significant when *p*<0.05 or, when using multiple hypothesis correction, adjusted-*p* <0.05 or FDR<0.05. We used Deming regressions to compare variables that were measured with error (e.g., log_2_ fold-change per Xi vs log_2_ fold-change per Chr Y). We calculated Deming regressions using the R package “deming” v1.4. For weighted Deming regressions, we included error values using the “xstd” and “ystd” arguments.

### KEY RESOURCES TABLE

#### Supplemental Table Legends

Table S1. Description of euploid and aneuploid donors.

Table S2. NPX and PAR gene linear regressions to Xi or Chr Y dosage.

Table S3. NPY gene linear regressions to Xi or Chr Y dosage.

Table S4. Autosomal linear regressions to Xi or Chr Y dosage.

Table S5. Autosomal linear regressions to Xi or Chr Y dosage, controlling for age.

Table S6. Gene set enrichment results for CD4+ T cells.

Table S7. Gene set enrichment results for monocytes.

Table S8. Donor samples from *in vitro* CD4+ T cell activation.

Table S9. Autosomal linear regressions to Xi of *in vitro* unstimulated and stimulated CD4+ T cells.

Table S10. Gene set enrichment results for *in vitro* unstimulated and stimulated CD4+ T cells.

Table S11. Deming regression and Pearson correlation stastistics between response to Xi vs. Chr Y of Hallmark functional pathways (restricted to leading edge genes).

