## Supplementary Figures for "Stable and robust Xi and Y transcriptomes drive cell-type-specific autosomal and Xa responses *in vivo* and *in vitro* in four human cell types"

#### Figure S1

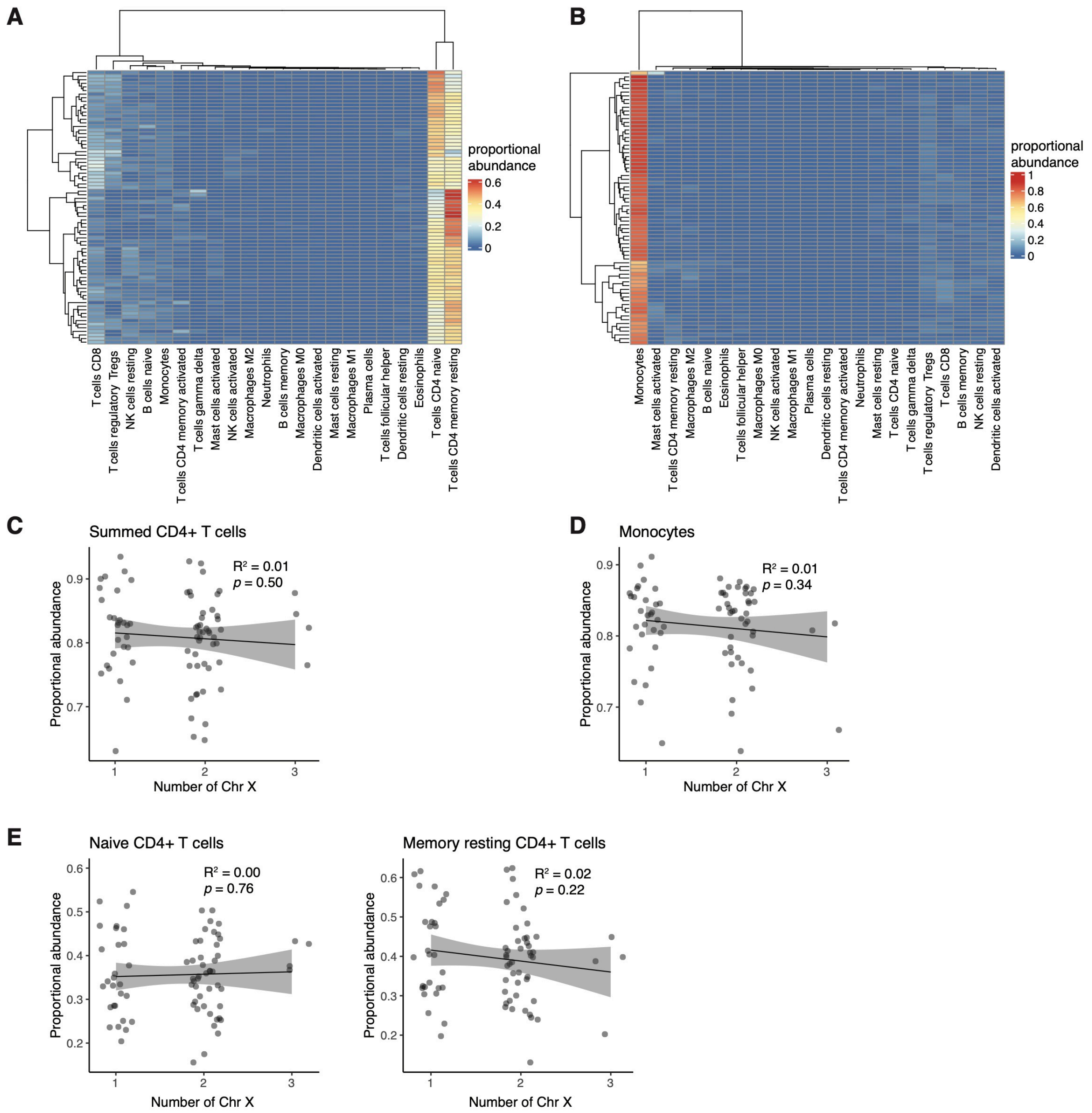


**Figure S1. Computational cell-type deconvolution confirms enrichment of CD4+ T cells and monocytes.**

**(A, B)** Heatmaps of estimated proportional abundances, as predicted by CIBERSORTx, for CD4+ T cell samples **(A)** and monocytes **(B)**. Each row is a single sample; columns indicate cell type identities. **(C)** Summed estimated proportional abundances of CD4+ T cells within a given sample, plotted by donor Chr X count. Linear regression statistics shown, indicating no significant change in CD4+ T cell abundance with the number of Chr X. **(D)** Estimated proportional abundances of monocyte samples, plotted by donor Chr X count. Linear regression statistics shown, indicating no significant change in monocyte abundance with the number of Chr X. **(E)** Estimated proportional abundances of naïve CD4+ T cells and memory resting CD4+ T cells. Linear regression statistics shown, indicating no significant change in CD4+ T cell abundance by Chr X count.

#### Figure S2

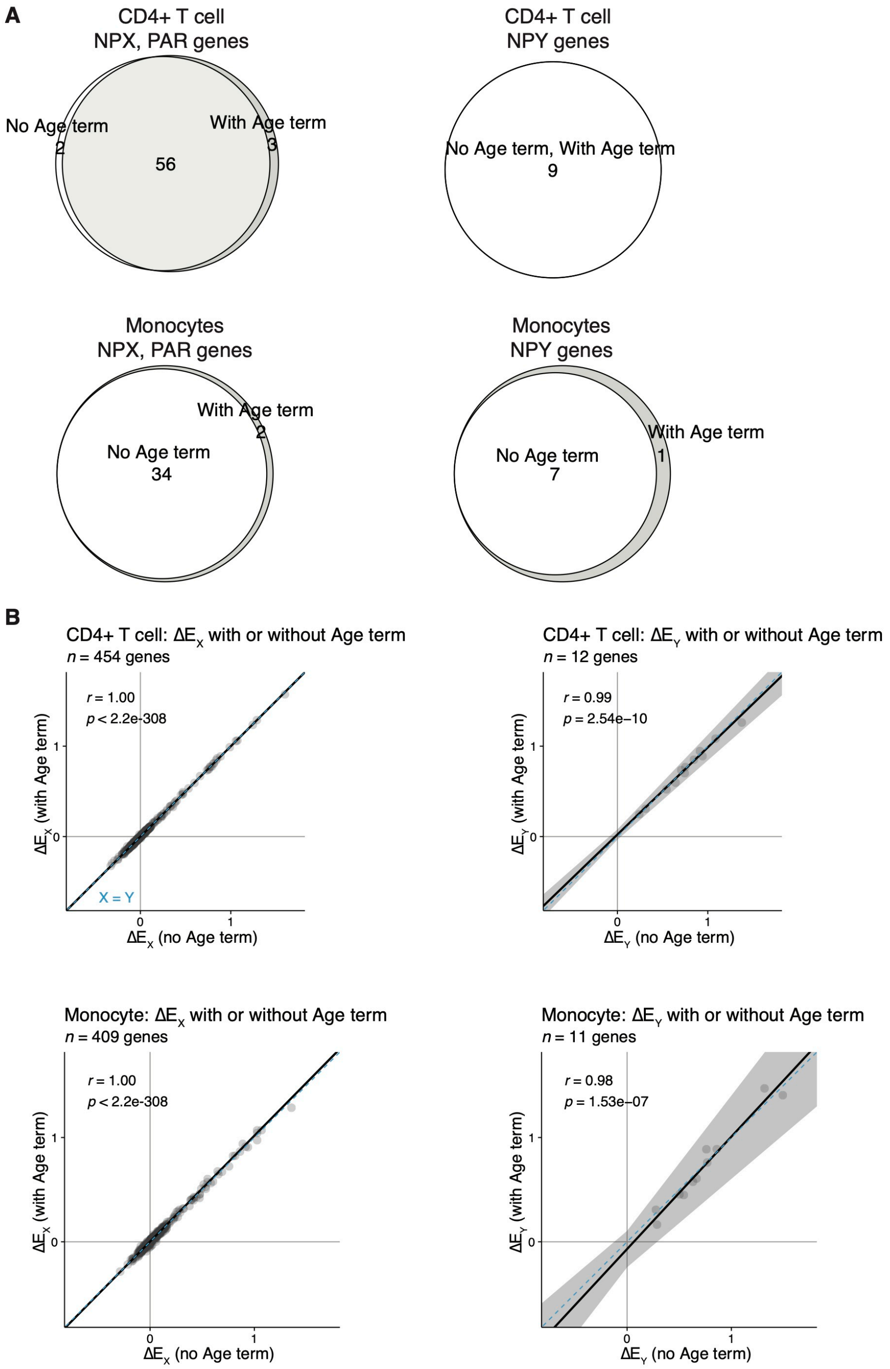


**Figure S2. Inclusion of age in linear model does not impact $\Delta E_X$ or $\Delta E_Y$.**

**(A)** Venn diagrams representing the number of X- or Y-chromosomal genes with significant responses to Chr X or Chr Y dosage, as indicated, with or without inclusion of age as a variable in the linear model. **(B)** Scatter plots of $\Delta E_X$ or $\Delta E_Y$ values, as indicated, in CD4+ T cells and monocytes with or without inclusion of age as a variable in the linear model. Deming regression line in black with 95% confidence intervals shaded gray; blue dotted lines indicate X=Y identity line. Pearson correlation statistics are shown.

#### Figure S3

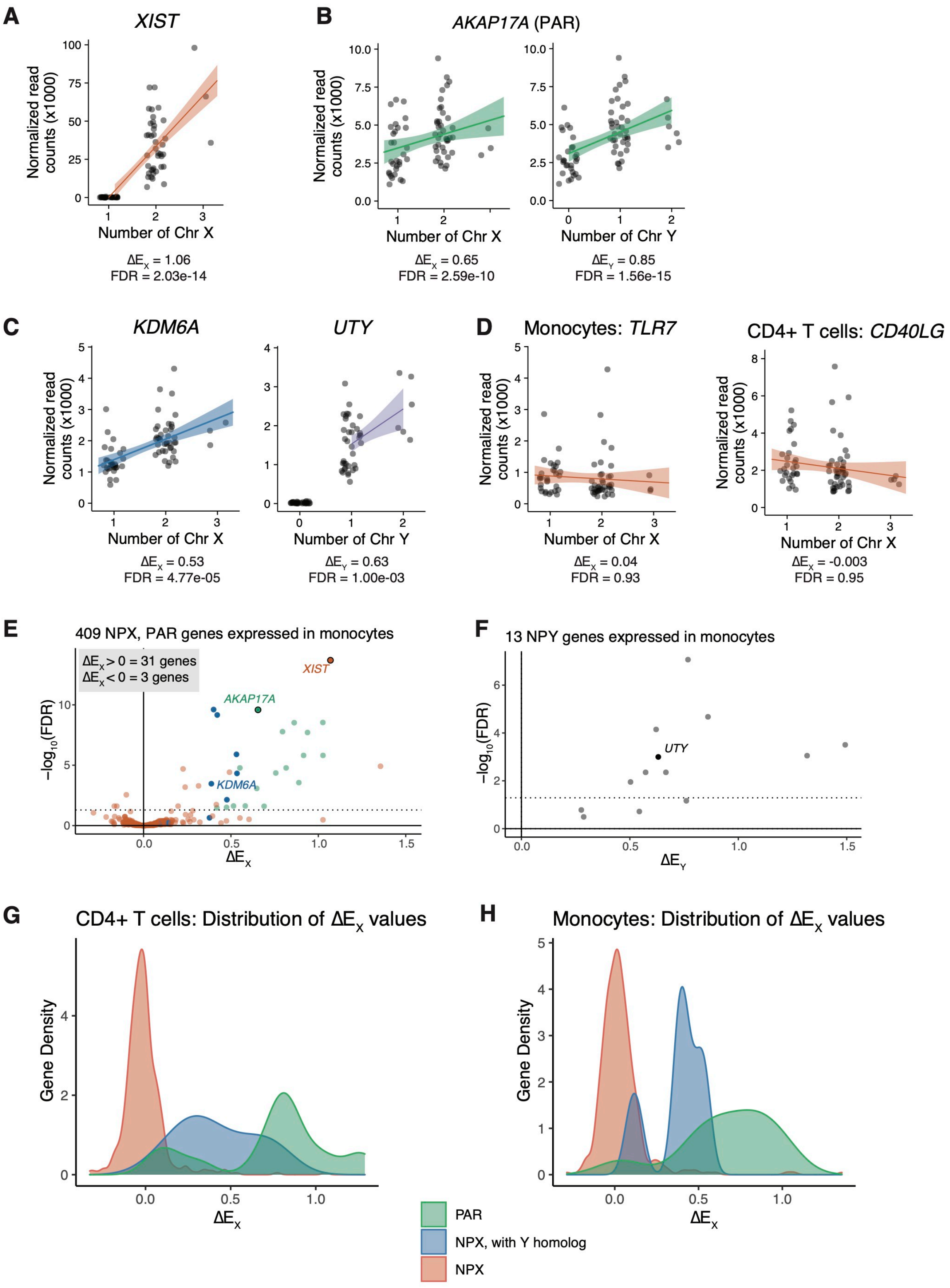

A
XIST
Normalized read counts (x1000)
Number of Chr X
ΔE_X = 1.06
FDR = 2.03e-14
B
AKAP17A (PAR)
Number of Chr X
ΔE_X = 0.65
FDR = 2.59e-10
Number of Chr Y
ΔE_Y = 0.85
FDR = 1.56e-15
C
KDM6A
Number of Chr X
ΔE_X = 0.53
FDR = 4.77e-05
UTY
Number of Chr Y
ΔE_Y = 0.63
FDR = 1.00e-03
D
Monocytes: TLR7
Number of Chr X
ΔE_X = 0.04
FDR = 0.93
CD4+ T cells: CD40LG
Number of Chr X
ΔE_X = -0.003
FDR = 0.95
E
409 NPX, PAR genes expressed in monocytes
ΔE_X > 0 = 31 genes
ΔE_X < 0 = 3 genes
XIST
AKAP17A
KDM6A
-log10(FDR)
ΔE_X
F
13 NPY genes expressed in monocytes
UTY
-log10(FDR)
ΔE_Y
G
CD4+ T cells: Distribution of ΔE_X values
Gene Density
ΔE_X
H
Monocytes: Distribution of ΔE_X values
Gene Density
ΔE_X
PAR
NPX, with Y homolog
NPX

**Figure S3. Expression of X- and Y-chromosomal genes respond to Xi and Y dosage in monocytes.**

**(A - D)** Normalized read counts (x1000) by sex chromosome dosage for *XIST* **(A)**, the PAR gene *AKAP17A* **(B)**, the NPX-NPY homologous pair *KDM6A* and *UTY* **(C)**, or the immune-related genes *TLR7* (in monocytes) and *CD40LG* (in CD4+ T cells) **(D)**. Regression lines with confidence intervals are shown. **(E, F)** Volcano plots of $\Delta E_X$ values of all expressed NPX and PAR genes **(E)** or $\Delta E_Y$ values of expressed NPY genes **(F)** in monocytes. Dotted horizontal lines indicate FDR = 0.05. In **(A)**, genes are annotated by gene class: PAR genes in green, NPX genes with NPY homologs in blue, and NPX genes with no NPY homolog in orange. **(G, H)** Density plots showing distributions of $\Delta E_X$ values by gene class in CD4+ T cells **(G)** and monocytes **(H)**.

**Figure S4**

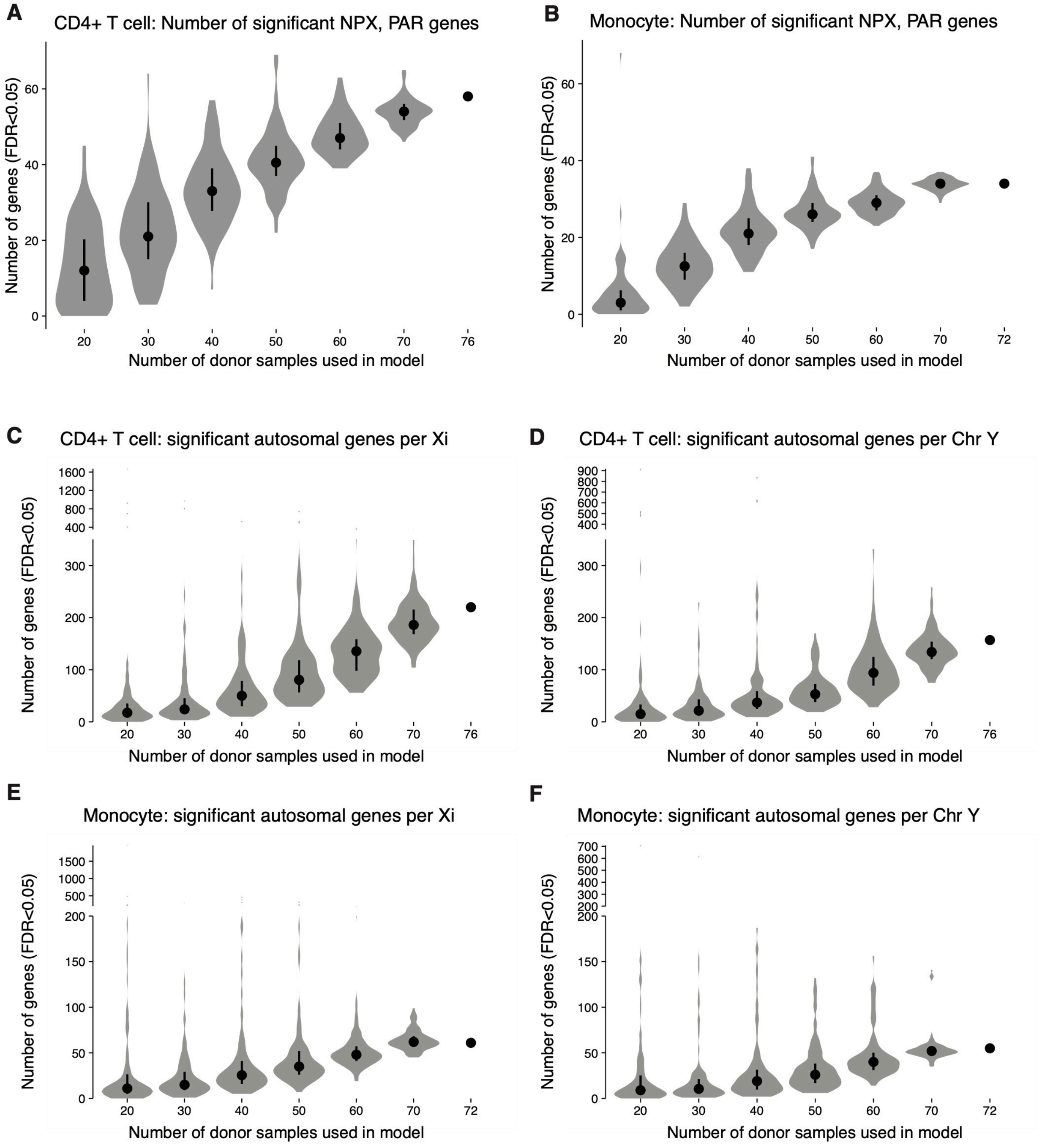


**Figure S4. Saturation analyses of detected significant gene responses to Xi or Chr Y dosage.** Subsampling the full dataset demonstrates that the number of significantly responsive NPX and PAR genes **(A, B)** as well as the number of autosomal genes with significant responses to Xi **(C,**

**E)** or Chr Y **(D, F)** dosage increase when more donor samples of CD4+ T cells and monocytes are utilized in the linear models. Violin plots show distributions of numbers of significant genes in 100 subsampling experiments.

### Figure S5

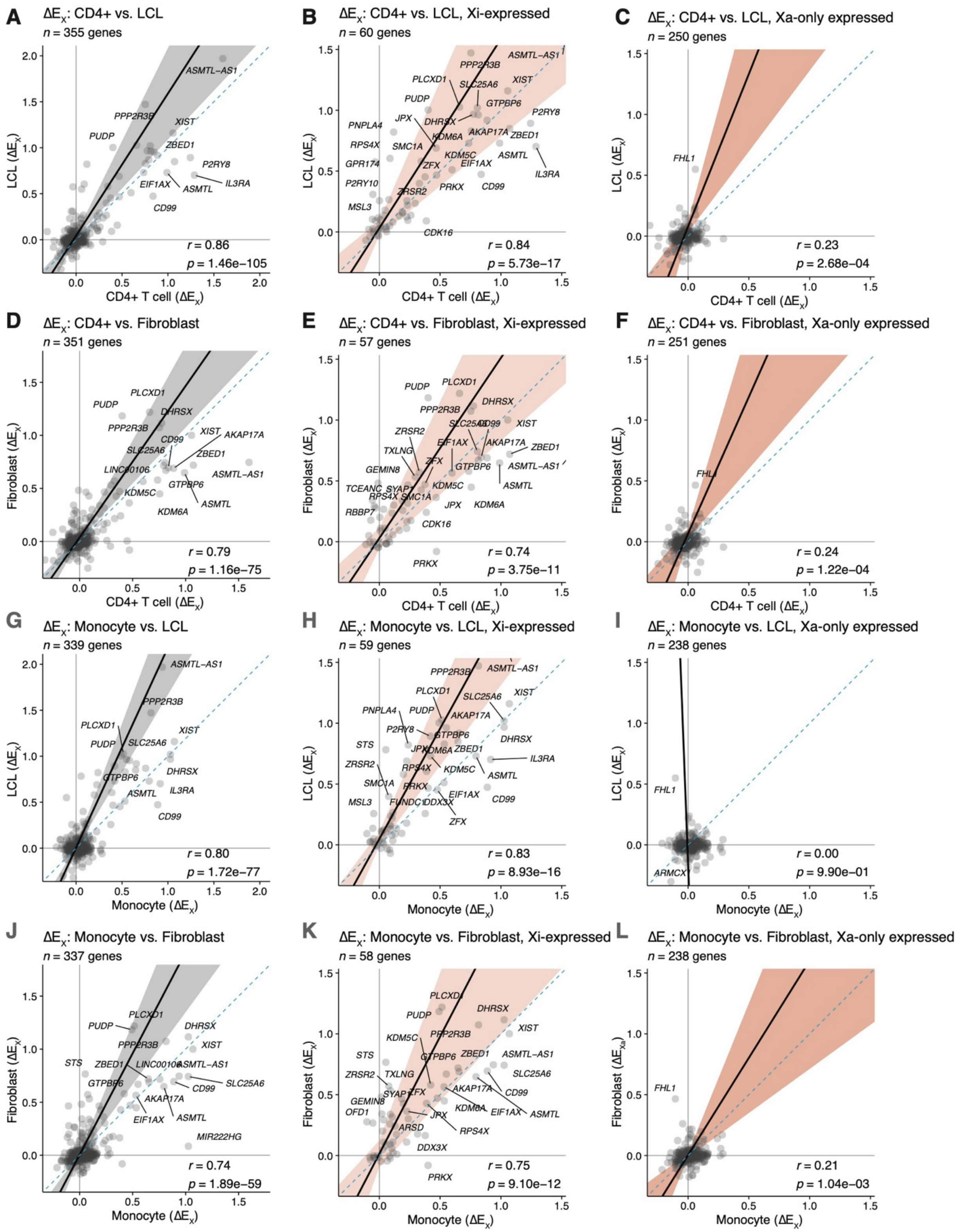

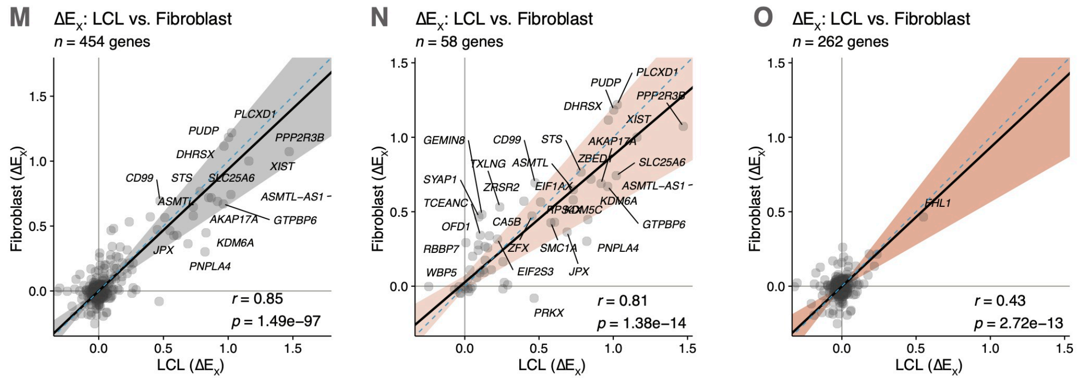


**Figure S5. X-chromosomal responses to Xi dosage are consistent across cell types, gene-by-gene.**

**(A - J)** Scatter plots comparing $\Delta E_X$ values for all NPX and PAR genes **(A, D, G, J, M)**, genes expressed on Xi, **(B, E, H, K, N)**, or genes only expressed on Xa **(C, F, J, L, O)** between the indicated cell types. The plotted genes are expressed in both of the indicated cell types. Deming regression line in black with 95% confidence intervals shaded gray; blue dotted lines indicate X=Y identity line. Pearson correlation statistics are shown. Confidence intervals are not plotted for **(I)**, as the intervals were broader than the plot axes.

**Figure S6**

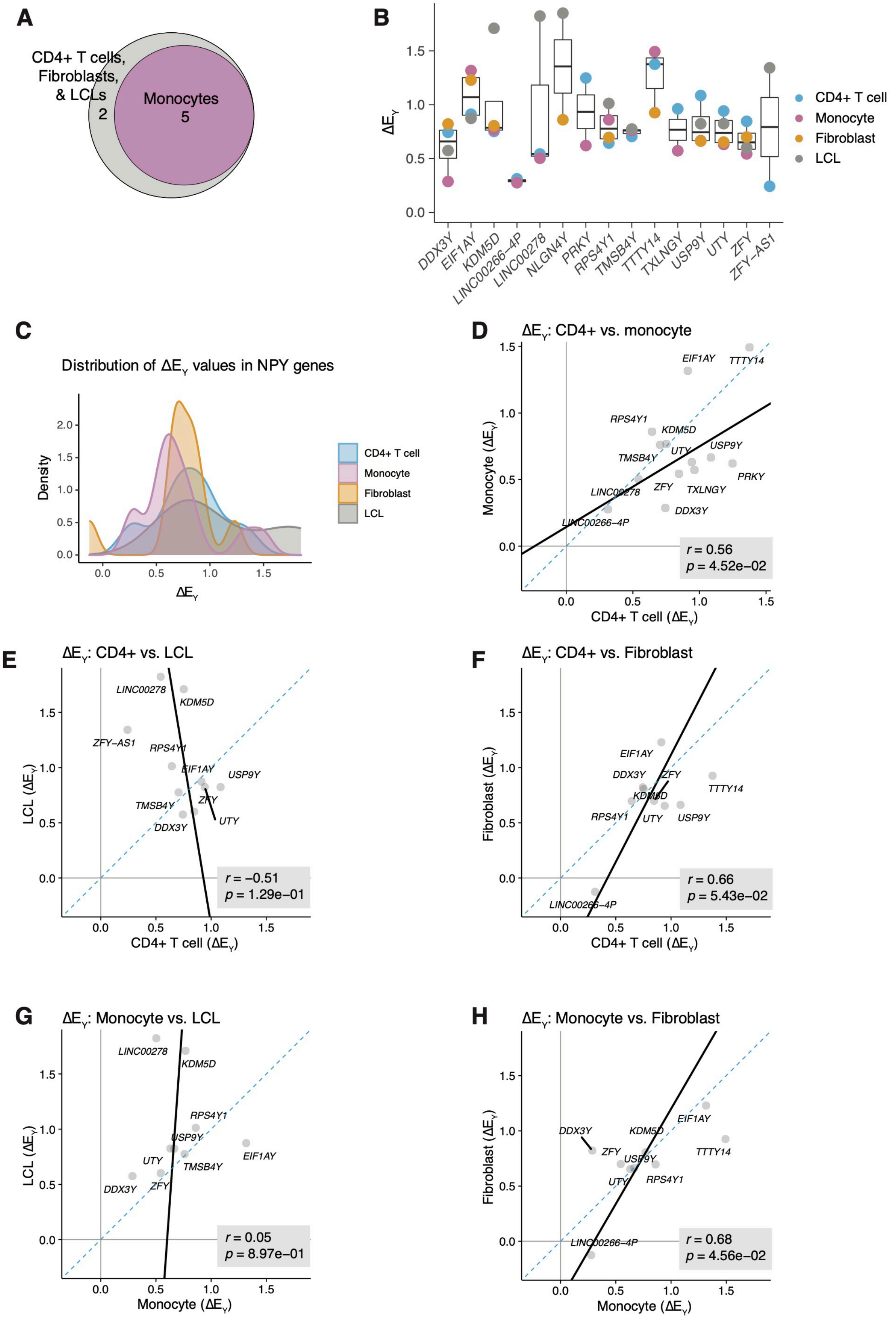

A
CD4+ T cells, Fibroblasts, & LCLs
2
Monocytes
5
B
ΔE_Y
1.5
1.0
0.5
0.0
DDX3Y
EIF1AY
KDM5D
LINC00266–4P
LINC00278
NLGN4Y
PRKY
RPS4Y1
TMSB4Y
TTTY14
TXLNGY
USP9Y
UTY
ZFY
ZFY–AS1
CD4+ T cell
Monocyte
Fibroblast
LCL
C
Distribution of ΔE_Y values in NPY genes
Density
ΔE_Y
CD4+ T cell
Monocyte
Fibroblast
LCL
D
ΔE_Y: CD4+ vs. monocyte
Monocyte (ΔE_Y)
CD4+ T cell (ΔE_Y)
r = 0.56
p = 4.52e−02
E
ΔE_Y: CD4+ vs. LCL
LCL (ΔE_Y)
CD4+ T cell (ΔE_Y)
r = −0.51
p = 1.29e−01
F
ΔE_Y: CD4+ vs. Fibroblast
Fibroblast (ΔE_Y)
CD4+ T cell (ΔE_Y)
r = 0.66
p = 5.43e−02
G
ΔE_Y: Monocyte vs. LCL
LCL (ΔE_Y)
Monocyte (ΔE_Y)
r = 0.05
p = 8.97e−01
H
ΔE_Y: Monocyte vs. Fibroblast
Fibroblast (ΔE_Y)
Monocyte (ΔE_Y)
r = 0.68
p = 4.56e−02

**Figure S6. NPY responses to Chr Y dosage across *in vitro* and *in vivo* cell types.**

**(A)** Venn diagram of NPY genes with statistically significant $\Delta E_Y$ values across the four cell types. **(B)** $\Delta E_Y$ values for each expressed NPY gene, colored by cell type; dots are only shown for genes expressed in the given cell type. **(C)** Density plot showing distribution of $\Delta E_Y$ values across the four cell types. **(D-H)** Scatter plots comparing $\Delta E_Y$ values between the indicated cell types. Deming regression line in black; blue dotted lines indicate X=Y identity line. Pearson correlation statistics are shown.

#### Figure S7

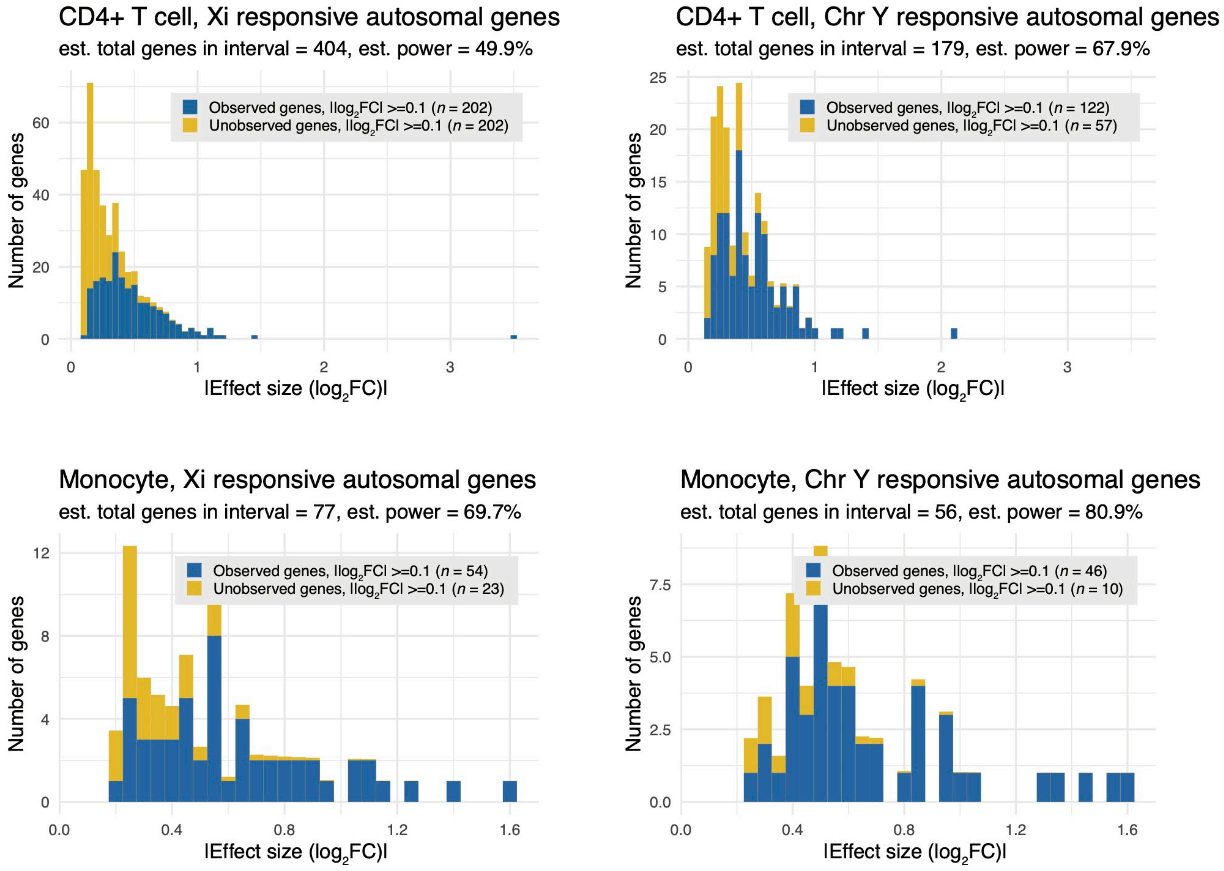


**Figure S7. Power analyses of observed and unobserved significant gene responses to Xi or Chr Y dosage in CD4+ T cells and monocytes.**

**(A – D)** Histograms depicting numbers of observed significantly responsive genes (in blue) and estimated unobserved significantly responsive genes (in yellow) by absolute effect size (|log2FC|) for CD4+ T cells' response to Xi **(A)** and Chr Y **(B)** and monocytes' response to Xi **(C)** and Chr Y **(D)**. Estimated numbers of unobserved genes were calculated via power analyses as described in Methods.

#### Figure S8

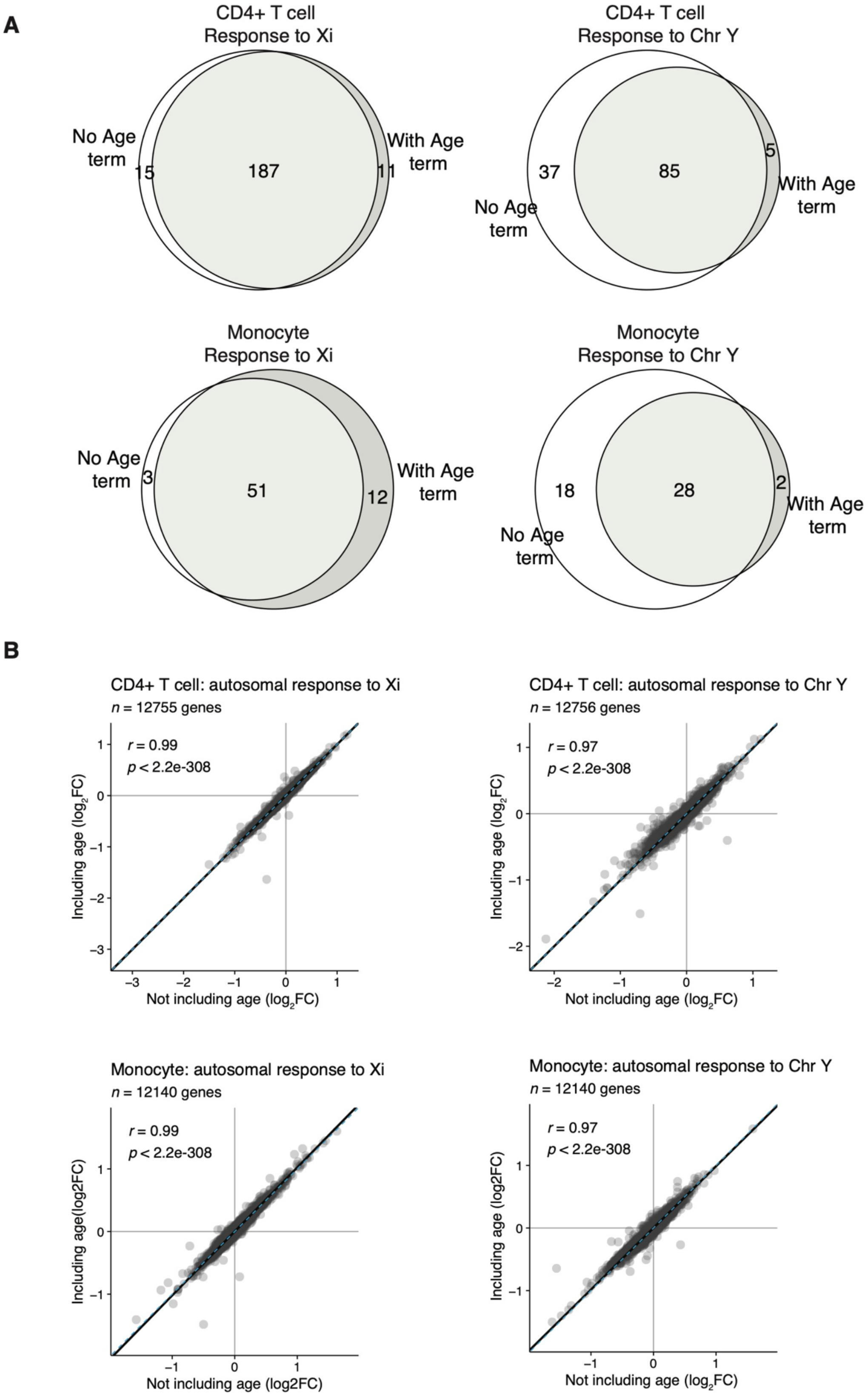
A
CD4+ T cell
Response to Xi
No Age term
15
187
11
With Age term
CD4+ T cell
Response to Chr Y
37
85
5
No Age term
With Age term
Monocyte
Response to Xi
No Age term
3
51
12
With Age term
Monocyte
Response to Chr Y
18
28
2
With Age term
No Age term
B
CD4+ T cell: autosomal response to Xi
n = 12755 genes
r = 0.99
p < 2.2e-308
Including age (log2FC)
Not including age (log2FC)
CD4+ T cell: autosomal response to Chr Y
n = 12756 genes
r = 0.97
p < 2.2e-308
Including age (log2FC)
Not including age (log2FC)
Monocyte: autosomal response to Xi
n = 12140 genes
r = 0.99
p < 2.2e-308
Including age(log2FC)
Not including age (log2FC)
Monocyte: autosomal response to Chr Y
n = 12140 genes
r = 0.97
p < 2.2e-308
Including age (log2FC)
Not including age (log2FC)

**Figure S8. Inclusion of age in linear model does not alter estimates of autosomal response to Xi or Chr Y dosage.**

**(A)** Venn diagrams representing the numbers of autosomal genes with significant responses to Xi or Chr Y dosage, as indicated, with or without inclusion of age as a variable in the linear model.

**(B)** Scatter plots of $\log_2$ fold-changes per Xi or Chr Y, as indicated, in CD4+ T cells and monocytes with or without inclusion of age as a variable in the linear model. Deming regression line in black with 95% confidence intervals shaded gray; blue dotted lines indicate X=Y identity line. Pearson correlation statistics are shown.

#### Figure S9

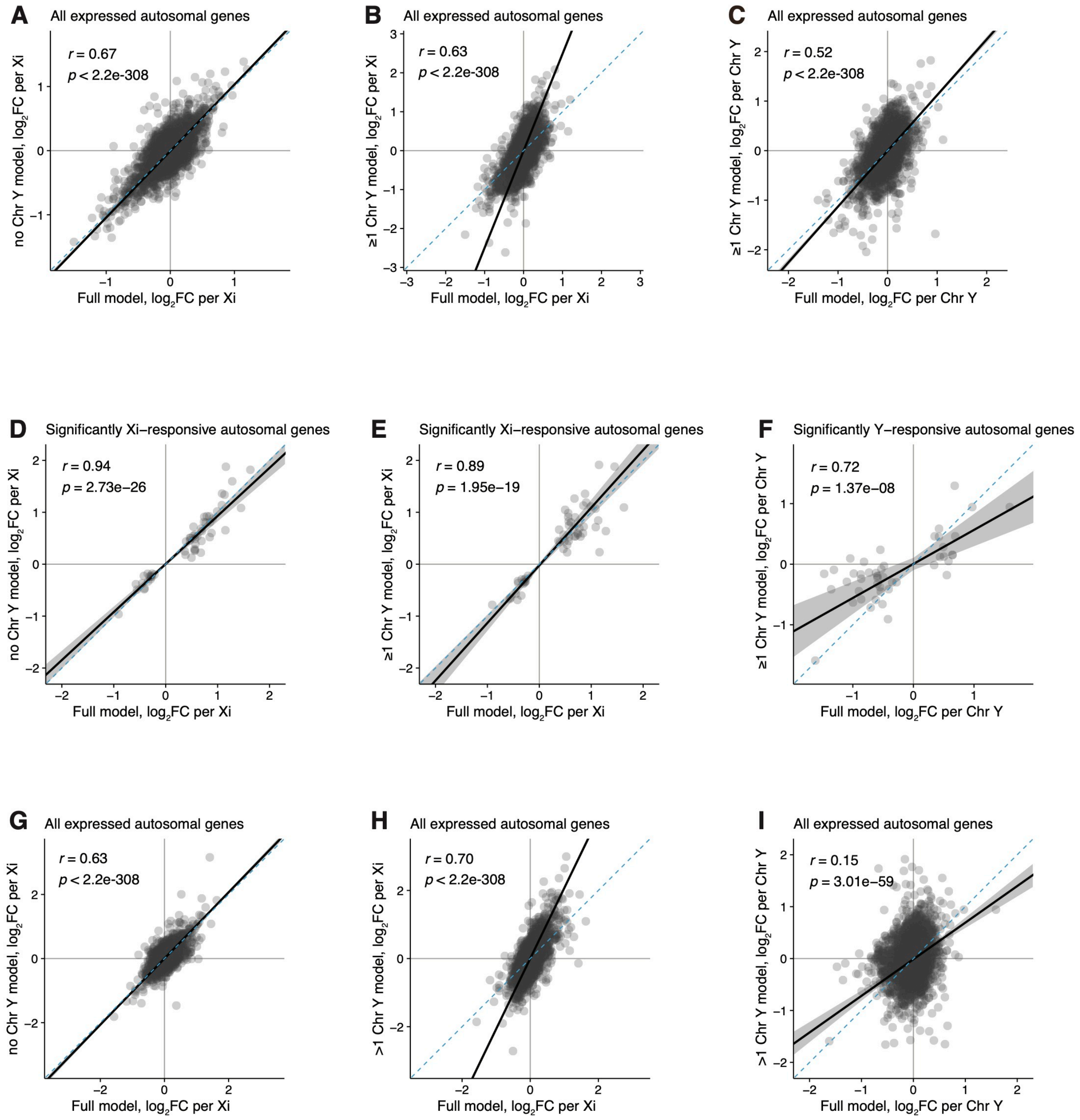


**Figure S9. Linear models built with no Chr Y or at least one Chr Y correlate with the full model containing all karyotypes.**

**(A)** Scatter plot of $\log_2$ fold-changes per Xi for all expressed autosomal genes in CD4+ T cells, comparing effect sizes found in full model with all available karyotypes versus ‘no Chr Y’ subset model. **(B, C)** Scatter plot of $log_2$ fold-changes per Xi **(B)** or Chr Y **(C)** for all expressed autosomal genes in CD4+ T cells, comparing effect sizes from full model with all available karyotypes versus $log_2$ fold-changes in '≥1 Chr Y' subset model. **(D)** Scatter plot of $log_2$ fold-changes per Xi for genes with significant Xi response in monocytes, comparing effect sizes found in full model with all available karyotypes versus 'no Chr Y' subset model. **(E, F)** Scatter plot of $log_2$ fold-changes per Xi **(E)** or Chr Y **(F)** for significantly responsive autosomal genes in monocytes, comparing effect sizes from full model with all available karyotypes versus $log_2$ fold-changes in '≥1 Chr Y' subset model. **(G)** Scatter plot of $log_2$ fold-changes per Xi for all expressed autosomal genes in monocytes, comparing effect sizes found in full model with all available karyotypes versus 'no Chr Y' subset model. **(H, I)** Scatter plot of $log_2$ fold-changes per Xi **(H)** or Chr Y **(I)** for all expressed autosomal genes in monocytes, comparing effect sizes from full model with all available karyotypes versus $log_2$ fold-changes in '≥1 Chr Y' subset model.

**Figure S10**

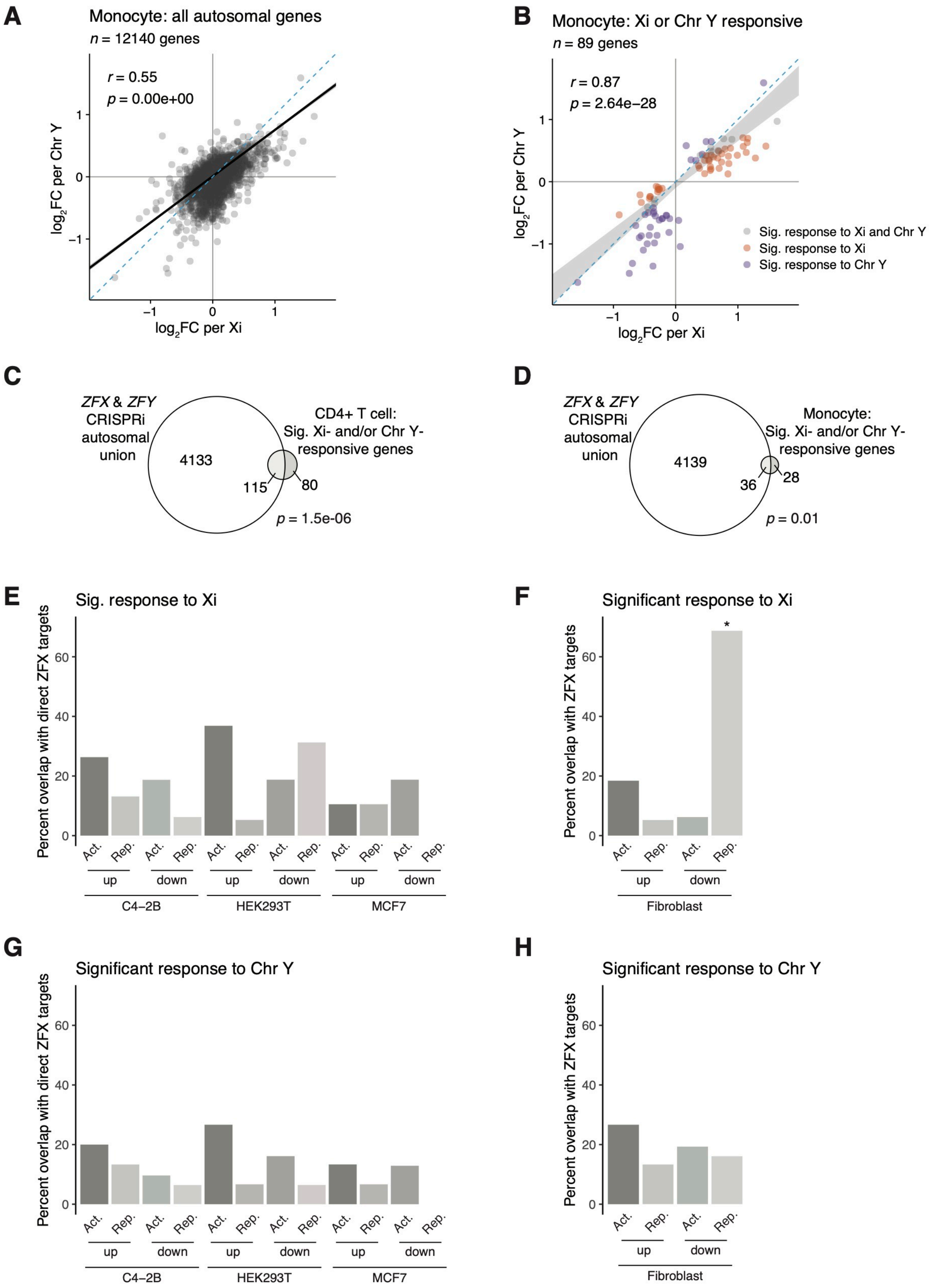
A
Monocyte: all autosomal genes
n = 12140 genes
r = 0.55
p = 0.00e+00
log2FC per Chr Y
log2FC per Xi
B
Monocyte: Xi or Chr Y responsive
n = 89 genes
r = 0.87
p = 2.64e−28
Sig. response to Xi and Chr Y
Sig. response to Xi
Sig. response to Chr Y
C
ZFX & ZFY CRISPRi autosomal union
4133
115
80
CD4+ T cell: Sig. Xi- and/or Chr Y-responsive genes
p = 1.5e-06
D
ZFX & ZFY CRISPRi autosomal union
4139
36
28
Monocyte: Sig. Xi- and/or Chr Y-responsive genes
p = 0.01
E
Sig. response to Xi
Percent overlap with direct ZFX targets
Act.
Rep.
up
down
C4−2B
HEK293T
MCF7
F
Significant response to Xi
Percent overlap with ZFX targets
Fibroblast
G
Significant response to Chr Y
Percent overlap with direct ZFX targets
H
Significant response to Chr Y
Percent overlap with ZFX targets

**Figure S10. ZFX and ZFY responsive genes explain a significant portion of the autosomal response to Xi dosage.**

**(A, B)** Scatter plot of $\log_2$ fold-changes per Xi versus Chr Y dosage of all expressed autosomal genes in monocytes **(A)** or limited to genes significantly responsive to Xi (orange), Chr Y (purple), or both (gray) **(B)**. (**C, D**) Venn diagrams displaying intersection of autosomal genes that respond to Xi or Chr Y in CD4+ T cells (**C**) or monocytes (**D**) with genes that respond to *ZFX* or *ZFY* knockdown via CRISPRi in fibroblasts. Autosomal gene sets were restricted to genes expressed in both fibroblasts and the corresponding immune cell type. **(E - H)** Bar plots showing percentage of genes responsive to Xi and/or Chr Y dosage in monocytes that were identified as ZFX direct target genes in MCF7, C4-2B, and HEK-293T cell lines, or ZFX responsive in fibroblasts. Genes are parsed by whether they significantly increased or decreased with Xi or Chr Y dosage ("up" or "down") and were activated or repressed by ZFX ("Act." or "Rep."). *p*-values reflect hypergeometric tests to identify significant enrichments of ZFX target genes in Chr X- or Y-responsive genes. Each comparison was restricted to genes expressed in both monocytes and the given *in vitro* cell line. Asterisks indicate the enrichment's *p*-value was lower than the Bonferroni-adjusted threshold of 0.001.

#### Figure S11

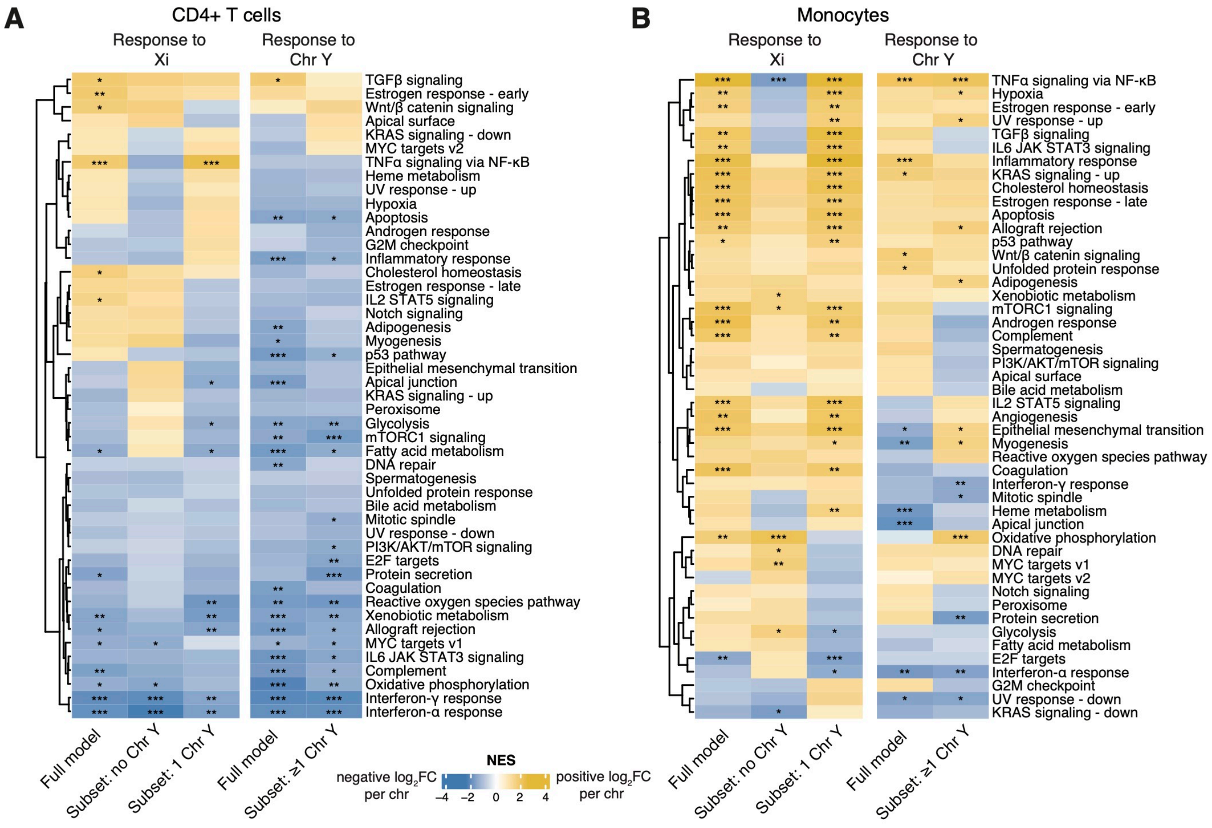


**Figure S11. CD4+ T cell functional enrichments remain largely consistent regardless of the presence or absence of Chr Y.**

**(A, B)** Heatmaps of normalized enrichment scores (NES) from gene set enrichment analysis with Hallmark gene sets in CD4+ T cells **(A)** and monocytes (**B**) for either the full models or subset models as indicated; significant enrichments indicated by asterisks (* = *p*-adj <0.05; ** = *p*-adj < 0.01; *** = *p*-adj < 0.001).

**Figure S12**

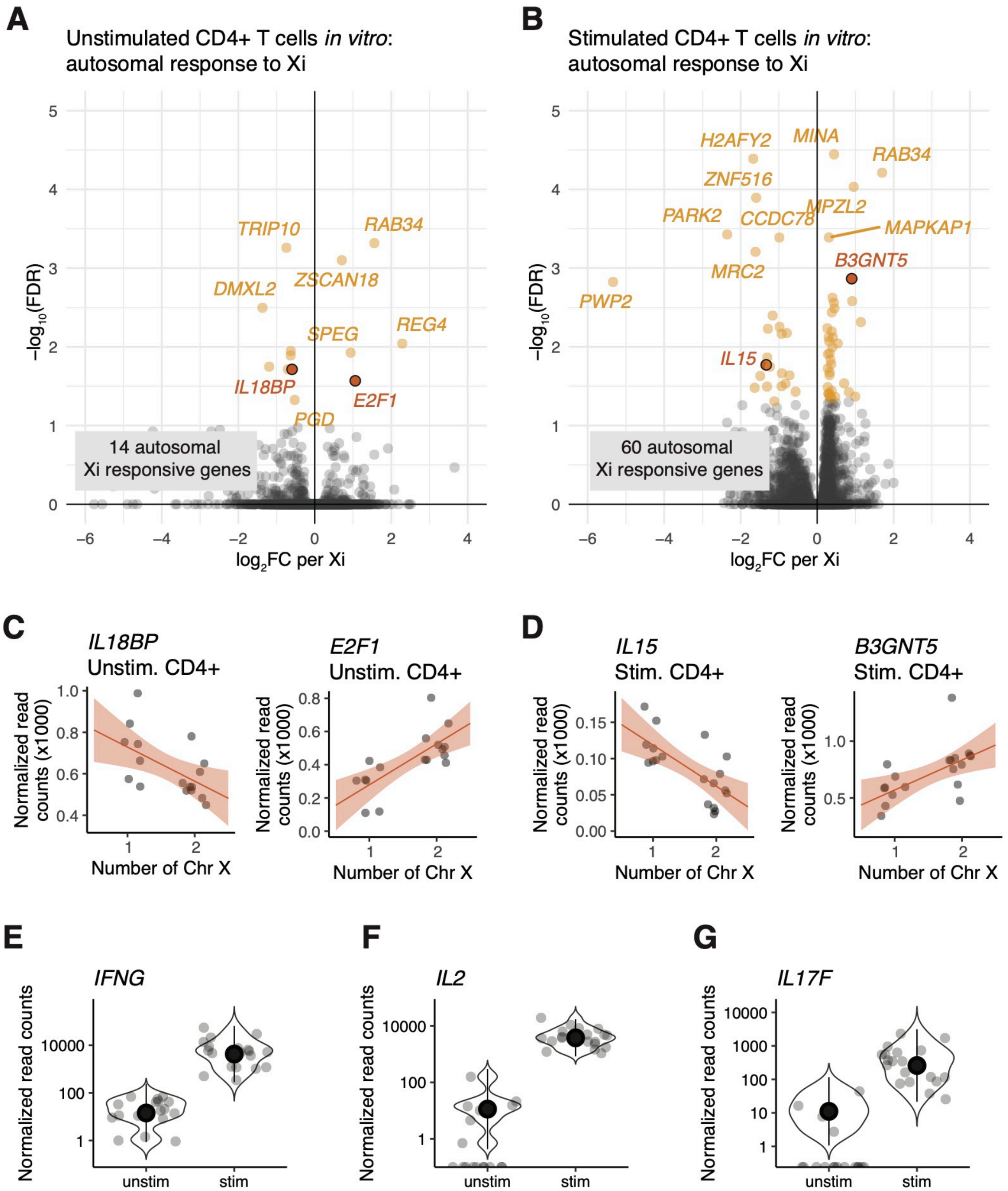


**Figure S12. Xi dosage has significant effects on CD4+ T cell activation.**

**(A, B)** Volcano plots of autosomal responses to Xi dosage for *in vitro* unstimulated CD4+ T cells (five positively and nine negatively responsive genes) **(A)** and *in vitro* stimulated CD4+ T cells

(37 positively and 23 negatively responsive genes) **(B)**. **(C, D)** Examples of positively and negatively Xi responsive genes in unstimulated CD4+ T cells **(C)** or stimulated CD4+ T cells **(D)**. Regression lines with confidence intervals are shown. **(E-G)** Normalized read counts showing increased *IFNG*, *IL2*, and *IL17F* gene expression with the addition of T cell activation beads, indicating that CD4+ T cell activation was successful.

### Figure S13

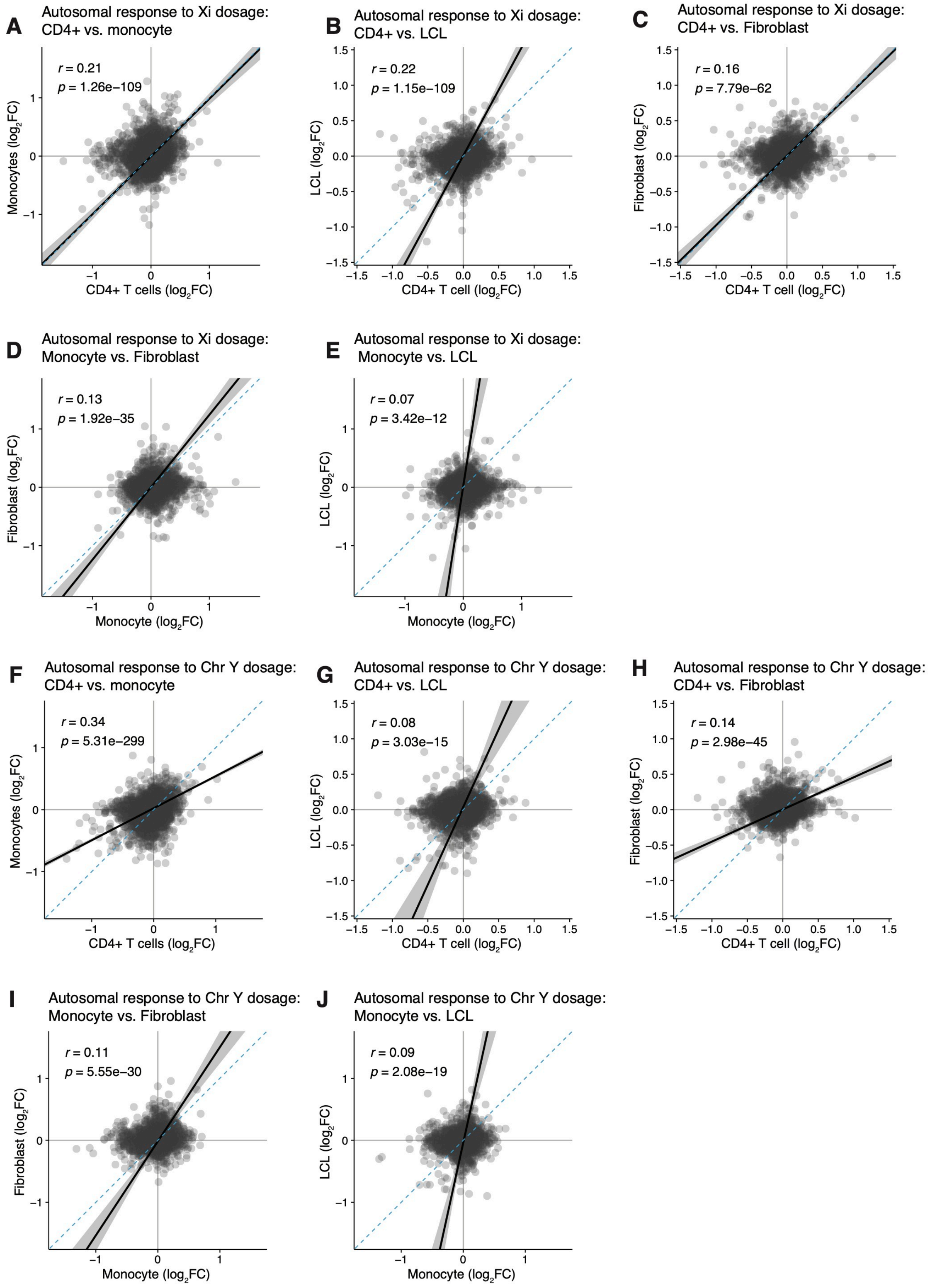
A
Autosomal response to Xi dosage:
CD4+ vs. monocyte
r = 0.21
p = 1.26e−109
Monocytes (log2FC)
CD4+ T cells (log2FC)
B
Autosomal response to Xi dosage:
CD4+ vs. LCL
r = 0.22
p = 1.15e−109
LCL (log2FC)
CD4+ T cell (log2FC)
C
Autosomal response to Xi dosage:
CD4+ vs. Fibroblast
r = 0.16
p = 7.79e−62
Fibroblast (log2FC)
CD4+ T cell (log2FC)
D
Autosomal response to Xi dosage:
Monocyte vs. Fibroblast
r = 0.13
p = 1.92e−35
Fibroblast (log2FC)
Monocyte (log2FC)
E
Autosomal response to Xi dosage:
Monocyte vs. LCL
r = 0.07
p = 3.42e−12
LCL (log2FC)
Monocyte (log2FC)
F
Autosomal response to Chr Y dosage:
CD4+ vs. monocyte
r = 0.34
p = 5.31e−299
Monocytes (log2FC)
CD4+ T cells (log2FC)
G
Autosomal response to Chr Y dosage:
CD4+ vs. LCL
r = 0.08
p = 3.03e−15
LCL (log2FC)
CD4+ T cell (log2FC)
H
Autosomal response to Chr Y dosage:
CD4+ vs. Fibroblast
r = 0.14
p = 2.98e−45
Fibroblast (log2FC)
CD4+ T cell (log2FC)
I
Autosomal response to Chr Y dosage:
Monocyte vs. Fibroblast
r = 0.11
p = 5.55e−30
Fibroblast (log2FC)
Monocyte (log2FC)
J
Autosomal response to Chr Y dosage:
Monocyte vs. LCL
r = 0.09
p = 2.08e−19
LCL (log2FC)
Monocyte (log2FC)

**Figure S13. Autosomal responses to Xi dosage are cell-type-specific.**

**(A – E)** Scatter plots of $log_2$ fold-changes for all expressed autosomal genes in response to Xi dosage between the indicated cell types. Deming regressions (solid black line with 95% confidence intervals shaded gray) and Pearson correlation statistics are shown. **(F – J)** Scatter plots of $log_2$ fold-changes for all expressed autosomal genes in response to Chr Y dosage between the indicated cell types. Deming regression line in black with 95% confidence intervals shaded gray; blue dotted lines indicate X=Y identity line. Pearson correlation statistics are shown.
